# Pitx2-associated early-onset glaucoma alters corneal innervation and sensory function in a sex-specific manner

**DOI:** 10.64898/2026.04.28.721459

**Authors:** Solange Sarkis, Luna Dumon, Mélissa Girard, Chantal Cazvieille, Marina Schverer, Vincent Daien, Bianca Johansen, Cécile Delettre, Chloé Chamard, Frederic Michon

**Affiliations:** Institute for Neurosciences of Montpellier, Univ Montpellier, INSERM, Montpellier, France; Laboratoires Théa, Clermont-Ferrand, France; Department of Ophthalmology, CHU Montpellier, Gui de Chauliac Hospital, Montpellier, France; The Save Sight Institute, Sydney Medical School, The University of Sydney, Sydney, NSW, Australia

**Author notes:** Contributed equally (Bianca Johansen, Cécile Delettre, Chloé Chamard, Frederic Michon).

**Keywords:** Early-onset glaucoma, Pitx2, Corneal neurobiology, Trigeminal ganglion, Sexual dimorphism, Transcriptomic

## Abstract

**Purpose:** Pitx2-associated developmental glaucoma is characterized by anterior segment dysgenesis, ocular hypertension, and optic neuropathy. Its consequences for corneal sensory innervation remain poorly understood. We investigated whether this disease alters corneal nerve structure and sensory function in a sex-dependent manner.

**Methods:** Male and female Pitx2^egl1/egl1^ and Pitx2^+/+^ mice were examined at 1 and 3 months. Ocular phenotyping included intraocular pressure, fundus imaging, visual evoked potentials, and optic nerve ultrastructure. RNA sequencing of corneas and trigeminal ganglia was performed at 3 months. Corneal innervation was assessed by βIII-tubulin immunofluorescence and volumetric quantification of nerve fibers. Corneal sensitivity was measured using Von Frey filaments.

**Results:** Pitx2^egl1/egl1^ mice developed progressive ocular hypertension, fundus abnormalities, reduced visual evoked potential amplitudes, and optic nerve degeneration, supporting the model as early-onset glaucoma. Baseline sex-related transcriptional differences were limited in both cornea and trigeminal ganglia. In contrast, Pitx2 mutation induced sex-dependent molecular responses. Female corneas showed broader transcriptional changes enriched in inflammatory, stress-response, and tissue-remodeling pathways, whereas male corneas showed a more restricted response involving metabolic and homeostatic processes. Similar sex divergence was observed in trigeminal ganglia. Corneal nerve fiber volume was reduced in both sexes at 3 months but not at 1 month, whereas reduced sensitivity was detected only in mutant males.

**Conclusions:** This study identifies sexual dimorphism as a component of Pitx2-associated developmental glaucoma. Furthermore, our findings suggest that glaucoma affects the corneal sensory system beyond optic nerve pathology, highlighting a potentially overlooked dimension of disease relevant to ocular surface monitoring and patient management.

**Highlights:**

- Disruption of the cornea–trigeminal ganglion axis with coordinated molecular and functional alterations in Pitx2-associated early-onset glaucoma
- Sex-dependent modifications in both cornea and trigeminal ganglion responses to early-onset glaucoma
- Progressive corneal neurodegeneration in early-onset glaucoma

## 1. Introduction

Mutations in Pitx2 are a well-established cause of early-onset developmental glaucoma, a severe ocular disorder characterized by anterior segment dysgenesis, including malformation of the trabecular meshwork, cornea, and iris [1–5]. Pitx2 encodes a paired-like homeodomain transcription factor that regulates multiple aspects of ocular morphogenesis, and its haploinsufficiency underlies Axenfeld-Rieger syndrome and related phenotypes, which often progress to glaucoma during infancy or adolescence [4,6,7]. While the structural consequences of Pitx2 deficiency have been well characterized, much less is known about its impact on the corneal nerve network, which plays a central role in ocular surface homeostasis and sensory function [5,6,8–10].

The cornea is one of the most densely innervated tissues in the human body, and its sensory nerves mediate protective reflexes, epithelial regeneration, and tear production [9–12]. A disruption in corneal innervation can lead to decreased corneal sensitivity, impaired epithelial regeneration, and increased susceptibility to injury or infection [11,13]. Although changes in the corneal nerve structure have been described in the context of various eye diseases, the impact of glaucoma on corneal innervation remains poorly understood [14–18]. This knowledge gap is particularly pronounced in developmental forms of glaucoma, where the effects on the corneal nerve structure and function have not been clearly defined. Considering that developmental glaucoma involves structural and functional alterations of the anterior segment [19–21], it is plausible that corneal innervation is secondarily affected, which could contribute to ocular surface disorders and long-term visual complications.

Emerging evidence indicates that sex can modulate ocular phenotypes in both developmental and adult-onset eye disorders [22–24]. Differences in hormone levels, gene expression, and epigenetic regulation have been implicated in sex-specific susceptibility and disease severity [25–27]. In animal models, sex differences influence corneal nerve density and regeneration after injury [28], suggesting that males and females may respond differently to genetic perturbations affecting ocular development. However, whether sex modifies corneal innervation or sensory outcomes in Pitx2-associated glaucoma has not been explored. Understanding such sex-specific effects is critical, as they may inform personalized approaches to monitoring and managing ocular surface health in patients with developmental glaucoma.

In this study, we aimed to investigate the effects of Pitx2-associated early-onset glaucoma on corneal innervation, sensory function, and molecular signatures, with attention to sex-specific differences. Using a combination of anatomical imaging, sensory assays, and transcriptomic profiling, we assessed corneal nerve architecture, functional responsiveness, and gene expression patterns in Pitx2^egl1/egl1^. We hypothesize that Pitx2 mutations disrupt corneal nerve structure and function in a sex-dependent manner and that these anatomical changes are accompanied by distinct transcriptional alterations.

## 2. Materials and Methods

## 2.1 Animals

Pitx2^egl1/+^ mice (C57BL/6J-Pitx2<^egl1^/Boc) were obtained from The Jackson Laboratory (Bar Harbor, ME, USA). Breeding was carried out in the animal facility (RAM-Neuro) of the Institute for Neurosciences of Montpellier to generate Pitx2^egl1/egl1^ and Pitx2^+/+^ mice. Genotyping was performed by the Montpellier Genomic Collection (MGC, Montpellier, France) using digital PCR with the following primers and probes: 59335 (common region (COM) forward: GAC TCA TTT CAC TAG CCA GCA G), 59336 (COM reverse: GAT TTC TTC GCG AGT GGA CA), 59337 Pitx2^+/+^ probe, FAM-labelled: CAG AGA AAC CGC TAC CCA GAC), and 59338 (Pitx2^egl1/egl1^ probe, HEX-labelled: CCA GAG AAA CCT CTA CCC AGA CA). Both male and female mice aged 1 and 3 months were used for the experiments.

All procedures were conducted in accordance with the European Union Directive 2010/63/EU on the protection of animals used for scientific purposes and the French regulations for animal experimentation (French Decree 2013-118). Experimental protocols were approved by the local ethical committee and authorized by the Ministère de la Recherche et de l’Enseignement Supérieur (authorization no. 2025062417478375, version 4). Mice were maintained under standard housing conditions (12 h light/12 h dark cycle, 21–22°C ambient temperature, 40–60% relative humidity) with food and water provided ad libitum.

### 2.2 Measurement of intraocular pressure (IOP)

Mice were anesthetized with isoflurane delivered via an induction chamber for 2 minutes prior to measurement. Following anesthesia, each mouse was positioned on an adjustable platform to align the eye at the same height as the tonometer, taking care to avoid any pressure on the head or neck. IOP was measured using iCare Tonolab rebound tonometer (Eickemeyer, Tuttlingen, Germany) in accordance with the manufacturer’s instructions, in a blinded manner. For each animal, six consecutive readings were taken, and the average value was used for analysis. All IOP measurements were performed between 2PM and 4 PM to minimize circadian variations.

### 2.3 Funduscopy

Twenty minutes before anesthesia, mydriatic agents (Mydriaticum, 2 mg/0.4 mL (Thea pharma, Clermont-Ferrand, France); and Neosynephrine 10%, Faure (Europhta, Monaco) were applied to achieve pupil dilation. Mice were then anesthetized with ketamine (80 mg/kg, Imalgene 1000) and xylazine (10 mg/kg, Rompun) administered intraperitoneally. The Ocry-Gel® (TVM) corneal ophthalmic gel was applied to the corneal surface before starting. Bright field filter was used and fundus images were acquired using a commercial camera and imaging system (Micron III; Phoenix Research Labs, Pleasanton, CA) in accordance with the manufacturer’s instructions. Still-frame images and video-rate sequences were acquired with Streampix III image acquisition software (NorPix, Montreal, Canada).

### 2.4 Visual evoked potential recordings

Electrophysiological recordings were performed in a darkroom in the morning, following overnight dark adaptation. On the day of the experiment, animals were anesthetized, and three electrodes were positioned as follows: the active electrode was placed at the nasal region, the reference electrode was placed at the base of the skull (over the occipital bone), and the ground electrode was inserted subcutaneously at the base of the tail. Visual evoked potentials were recorded using the Visio system (SIEM BioMédicale; Nîmes, France) following flash light stimulation. Recordings consisted of three phases of 60 flashes each, as previously described [29]. Flash duration was 5 ms, with a frequency of 1 Hz and an intensity of 159 cd·s·m⁻². Amplitudes and latencies obtained during each phase were averaged, and a cut-off filter was set at 35 Hz.

### 2.5 Ultrastructure analysis of the optic nerve by electron transmission microscopy

Optic nerves from female mice were dissected and immediately fixed overnight at 4°C in 0.1 M PHEM buffer (pH 7.2) containing 2.5% glutaraldehyde. Samples were then rinsed in PHEM buffer and post-fixed for 2 hours at room temperature in the dark with 0.5% osmium tetroxide and 0.8% potassium ferrocyanide. After two washes in distilled water, tissues were processed using a microwave tissue processor (Leica AMW) for resin embedding.

Tissues were dehydrated through a graded ethanol series (30–100%) followed by absolute acetone, infiltrated with epoxy resin (EMBed 812) via progressive acetone–resin baths, and polymerized at 60°C. Ultrathin sections (∼70 nm) were cut on a Leica-Reichert Ultracut E ultramicrotome and collected on copper grids at multiple levels of the optic nerve. Sections were contrasted with 1.5% uranyl acetate in 70% ethanol and lead citrate, then examined using a Tecnai F20 transmission electron microscope operated at 120 kV, equipped with a Veleta digital camera (Plateau de Microscopie Electronique, Institut des Neurosciences de Montpellier, INSERM U1298, Université Montpellier, Montpellier, France).

### 2.6 Corneal sensitivity test

The corneal sensitivity was measured using Von Frey filaments (Bioseb,Vitrolles, France; reference bio-VF-M). During testing, mice were gently immobilized, and filaments generating forces between 0.008 and 1 g were applied to the center of the cornea. The force was gradually increased by using filaments of higher weight until a blink reflex was observed. Each mouse was tested on three consecutive days, and the average value obtained from the three sessions was used for analysis. Corneal sensitivity was expressed as the inverse of the force required to elicit the blink reflex (g^-1^). All measurements were performed by the same experimenter under single-blinded conditions.

### 2.7 Sample collection and processing

Mice were euthanized by cervical dislocation. Eyes were collected by enucleation using curved scissors to sever the optic nerve and were briefly washed in PBS. Heads were then collected and immediately placed on ice. The skin was removed, and the skull was carefully opened. The trigeminal ganglia were subsequently dissected and isolated. For RNA-seq analysis, corneas and trigeminal ganglia were rapidly dissected under RNase-free conditions, snap-frozen in liquid nitrogen, and stored at −80 °C until further processing. Two corneas and trigeminal ganglia were pooled to generate each sample.

For immunofluorescence staining, eyes were fixed in 4% paraformaldehyde solution (Antigenfix) for 20 minutes and washed three times in PBS for 15 minutes each. Samples were then dehydrated for 2 hours in 50% ethanol/PBS and subsequently stored in 70% ethanol/PBS at 4 °C.

### 2.8 Immunofluorescence analysis of the corneas

Dissected and fixed corneas were blocked and permeabilized in a solution containing 2.5% fish skin gelatin (FSG; Sigma-Aldrich, G7765) and 5% goat serum (GS; Thermo Fisher Scientific, 16210064) in 0.5% Triton X-100 in PBS at room temperature. Tissues were subsequently incubated overnight at 4 °C with the primary antibody anti-βIII-tubulin (rabbit; Abcam, ab18207; 1:1000) prepared in 2.5% FSG and 5% GS in 0.1% Triton X-100 in PBS. After incubation, corneas were rinsed three times for 1 h each in 0.1% Triton X-100 in PBS at room temperature and then incubated with the secondary antibody Alexa Fluor 488–conjugated goat anti-rabbit IgG (H + L) (Thermo Fisher Scientific, A-11008; 1:500) under the same conditions and rinsed three times for 1 h each in 0,1% Triton X-100 in PBS at room temperature. Nuclei were counterstained with Hoechst 33342 (Thermo Fisher Scientific, H3570) for 10 min at room temperature, followed by a final rinse in PBS for 5 min.

Four radial incisions were made with a carbon steel surgical blade (15C, reference 0221, Swann-Morton) in each cornea to allow flat mounting. Corneas were then mounted in Vectashield antifade mounting medium (Vector Laboratories, H-1000) on glass slides, with the endothelial side facing downward toward the slide.

### 2.9 Imaging

Whole-cornea images were acquired as previously described [30] using a Leica Thunder Imager Tissue microscope equipped with the Navigator module and Large Volume Computational Clearing (LVCC) function. Image capture was performed with LAS X software (v3.7.4) using a 20×/0.55 objective. Image processing was conducted in Imaris (Bitplane, v10.0.0). Identical acquisition and processing parameters were applied to all samples within the same experimental set to ensure consistency.

### 2.10 Innervation volume measurement

All image analyses were performed by the same experimenter under single-blinded conditions. Using FIJI software (RRID:SCR_002285), a centred circular region corresponding to 50% of the corneal area was defined for each image. The circle radius was determined by measuring the corneal diameter twice with the measurement plug-in and calculating the mean value. A 50% circular crop of each cornea was then generated using the crop plug-in, followed by the clear outside plug-in to remove the surrounding area.

The resulting cropped images were converted to ims format using the Imaris Converter and analyzed in Imaris (Bitplane, version 10.0.0). Corneal nerve innervation density was quantified using the “Surface” tool, and the corresponding volume data (µm³) were extracted for statistical analysis. The same threshold settings were applied to all corneas within the same experimental cohort to ensure consistency in quantification.

### 2.11 RNA-Seq on corneas and trigeminal ganglia

The frozen corneas and trigeminal ganglia were shipped in dry ice to the Beijing Genomics Institute (BGI, Shenzhen, China) for library preparation and sequencing on the DNBSEQ platform. Total RNA was extracted using the RNeasy Mini Kit (Qiagen) according to the manufacturer’s instructions. Polyadenylated mRNA was enriched using oligo(dT)-coated magnetic beads and then fragmented. RNA quality and fragment size distribution were assessed using a Fragment Analyzer, and only samples meeting predefined quality control thresholds were selected for library preparation. Sequencing libraries were constructed using the BGI Optimal Dual-mode mRNA Library Prep Kit and sequenced on a DNBSEQ-T7 platform to generate 2 × 100 bp paired-end reads, yielding approximately 50 million clean reads per sample. Raw sequencing reads were processed with SOAPnuke to remove adapter sequences, low-quality bases, and contaminants. The filtered reads were then aligned to the mouse reference genome (Mus musculus, mm10, UCSC assembly v2201) using HISAT2. Gene-level read counts were quantified using HTSeq-count, producing the final raw count matrix for downstream analysis.

### 2.12 Gene Ontology analysis

To analyze the modulation of biological processes (BP) from the Gene Ontology (GO) database and the RNA-Seq data on corneas, statistical analyses were performed using R and RStudio (version 4.4.2). The BiocManager package from the Bioconductor open source software project was used (*clusterProfiler* and *AnnotationDbi* packages). Genes were then associated with biological processes and an overrepresentation (ORA) analysis was performed using the GO database, with the Mus musculus reference genome as the background dataset (org.Mm.eg.db). Results are presented as dot plots and show the top 20 upregulated or downregulated biological processes. KEGG (Kyoto Encyclopedia of Genes and Genomes) pathway enrichment analysis was conducted using R in combination with the Enrichr platform (via the enrichR package).

### 2.13 Statistical analysis

Data are presented as mean ± standard error of the mean (SEM). Statistical analyses were performed using GraphPad Prism (version 10.1.3, GraphPad Software, CA, USA). Comparisons between two groups were conducted using unpaired, two-tailed t-tests. Statistical significance was defined as *p < 0.05, **p < 0.01, ***p < 0.001, ****p < 0.0001.

For IOP measurement, immunofluorescence analysis of the corneas and corneal sensitivity, values from the left and right corneas of each animal were averaged to obtain a single data point.

## 3. Results

### 3.1 Validation of the Pitx2^egl1/egl1^ model of early onset glaucoma

Previous work has shown that Pitx2 mutations cause anterior segment defects associated with ocular hypertension and retinal ganglion cell loss [1,2,31]. Before examining the corneal consequences of this mutation, we first confirmed the extent to which Pitx2^egl1/egl1^ mice recapitulate the core features of early-onset glaucoma. To this end, we performed longitudinal analyses of IOP, fundus morphology, visual pathway function, and optic nerve integrity in male and female Pitx2^+/+^ and Pitx2^egl1/egl1^ mice at 1 and 3 months of age.

Across both time points, Pitx2^egl1/egl1^ mice presented elevated IOP compared with controls, with higher values observed at the earlier time point (21.36 ± 1.75 vs 18.06 ± 1.52 mmHg) and at 3 months of age (23.46 ± 2.84 vs 17.28 ± 1.78 mmHg), consistent with a progressive ocular hypertensive phenotype (Fig. 1A). This sustained increase in IOP indicates that the mutation induces early and persistent alterations in aqueous humour dynamics. While modest sex-specific differences were detected in mutants at later stages, with females exhibiting higher IOP than males (Fig. S1), these remained secondary to the overall hypertensive phenotype.

**Figure 1:**
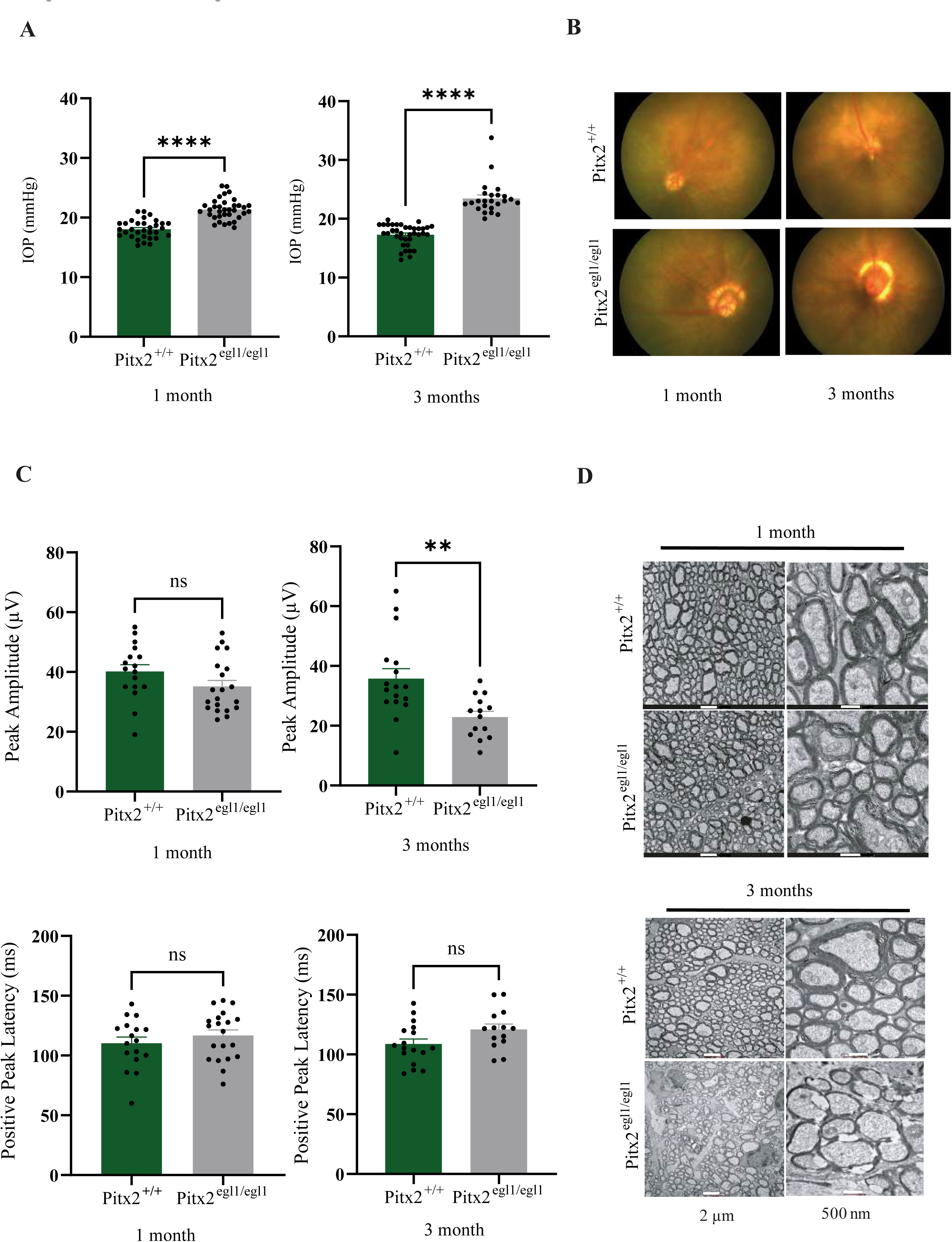
Ocular phenotyping of glaucoma-associated Pitx2 mutant mice across age and sex. (A) IOP measurements in male and female mice at 1 and 3 months of age. At 1 month: Pitx2^+/+^ (n = 32; 14 males) and Pitx2^egl1/egl1^ (n = 35; 24 males). At 3 months: Pitx2⁺/⁺ (n = 37; 16 males) and Pitx2 ^egl1/egl1^ (n = 24; 13 males). Data are presented in mmHg and bars represent mean IOP ± SEM, with individual data points shown for each genotype and sex. (B) Representative fundus images from Pitx2⁺/⁺ and Pitx2 ^egl1/egl1^ mice at 1 and 3 months of age. (C) Quantification of P1 latency (ms) and N1–P1 amplitude (µV) in male and female mice at 1 and 3 months of age. At 1 month: Pitx2⁺/⁺ (n = 17; 6 males, 11 females) and Pitx2 ^egl1/egl1^ (n = 14; 8 males, 6 females). At 3 months: Pitx2⁺/⁺ (n = 17; 6 males, 11 females) and Pitx2 ^egl1/egl1^ (n = 16; 6 males, 10 females). Data are presented as mean ± SEM with individual data points shown for each group. (D) Representative ultrastructural images of optic nerve cross-sections from Pitx2⁺/⁺ and Pitx2 ^egl1/egl1^ female mice at 1 and 3 months (scale bars: 2 µm and 500 nm). Statistical significance is indicated as ns (not significant), *p < 0.05, **p < 0.01, ***p < 0.001, ****p < 0.0001.

Fundus imaging further revealed progressive posterior segment abnormalities in Pitx2^egl1/egl1^ mice. The presence of a ring-like peripapillary structure surrounding the optic nerve head, which became more pronounced with age, is consistent with structural remodelling associated with glaucomatous damage (Fig. 1B). The absence of these features in control mice supports the specificity of this phenotype to the mutant condition.

To determine whether these alterations were associated with functional impairment of the visual pathway, visual evoked potentials were recorded. Recordings showed that Pitx2^egl1/egl1^ (22.93 ± 7.01 µV) exhibited significantly reduced peak amplitudes at 3 months compared to Pitx2^+/+^ mice (35.76 ± 13.66 µV), whereas no difference was observed at 1 month compared with controls (Fig. 1C). Positive-peak latencies were similar between mutants and controls at both ages, indicating that conduction timing along the visual pathway is preserved. In line with these functional deficits, ultrastructural analysis of the optic nerve revealed progressive degeneration in female mutant mice. While optic nerve organization was relatively preserved at 1 month, 3 months old Pitx2^egl1/egl1^ mice presented marked axonal disorganization, and myelin abnormalities, including thinning and focal delamination (Fig.1D), consistent with ongoing neurodegenerative processes.

Although some sex-related differences were detected for individual parameters, they did not alter the overall pattern of progressive glaucomatous damage and are therefore presented in the supplementary analyses (Fig. S2). Having validated the model at the ocular level, we next asked whether this pathology also extends to the corneal sensory axis, with particular attention to potential sex-dependent effects.

### 3.2 Progressive, sex-dependent disruption of the corneal sensory axis in Pitx2egl1/egl1 mice

We therefore examined corneal innervation and sensitivity over time in male and female mutant and control mice to determine when these alterations emerge and whether they differ by sex. At 1 month of age, corneal nerve structure (Fig. 2A) and sensitivity (Fig. 2C) were comparable between mutant and control mice in both sexes, indicating no detectable early deficits. However, by 3 months of age, progressive alterations became evident. Pitx2^egl1/egl1^ mice (male: 9.83 × 10^6^ µm^3^, female: (1.49 × 10^7^ µm^3^) showed reduced epithelial nerve density compared to control mice (male: 12.38 × 10^6^ µm^3,^ p = 0.0028, female: 1.01 × 10^7^ µm^3^, p= 0.0057), in both sexes. While control females had a slightly higher nerve volume than males, this normal sexual dimorphism was attenuated in the mutants (Fig. 2B). These results suggest that the Pitx2 gene mutation disrupts corneal nerve structure and alters normal sexual dimorphism.

**Figure 2:**
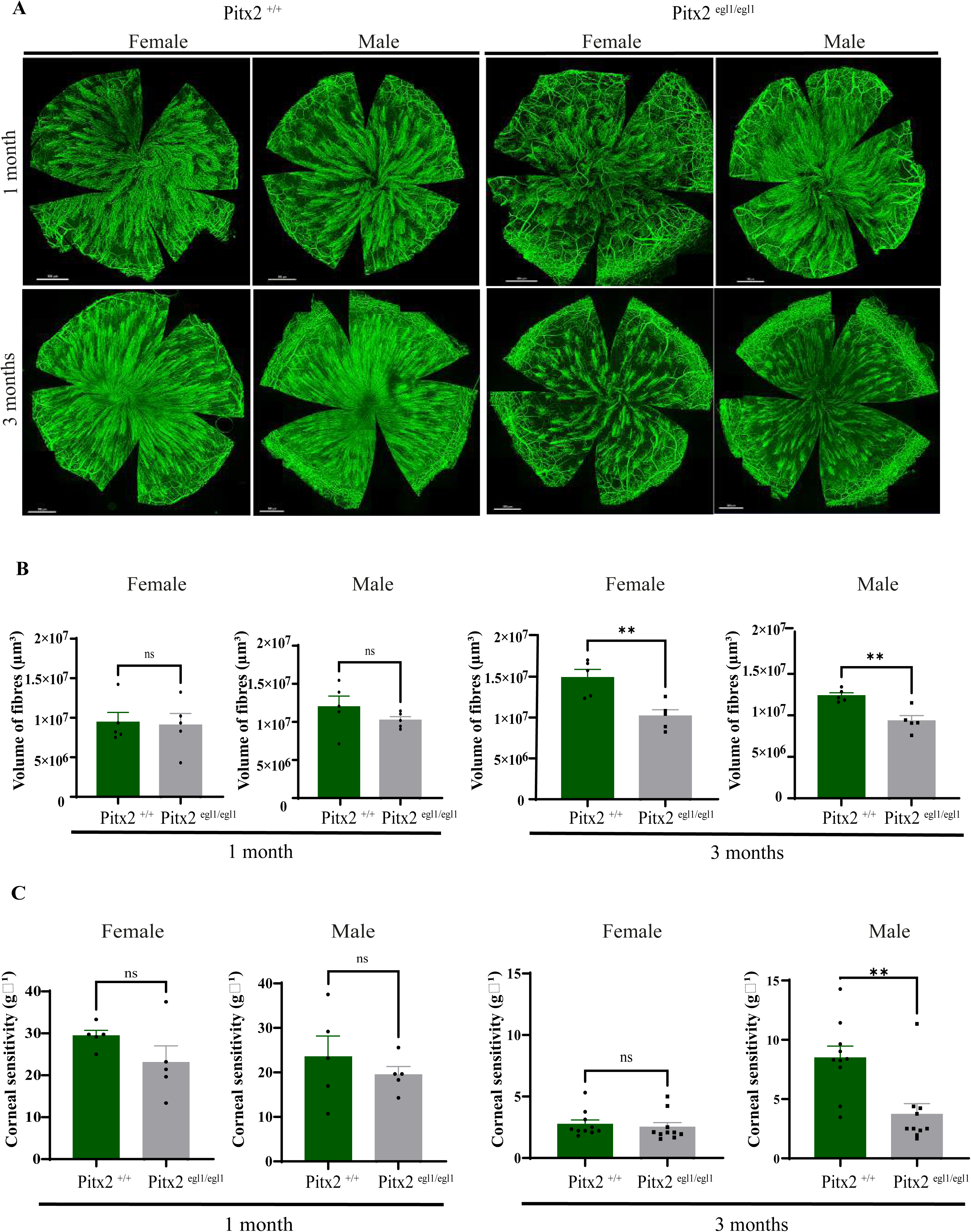
Corneal epithelial innervation and corneal sensitivity in Pitx2⁺/⁺ and Pitx2^egl1/egl1^ mice at 1 and 3 months. (A) Whole-mount images of the corneal subbasal epithelial nerve plexus from female and male Pitx2⁺/⁺ and Pitx2^egl1/egl1^ mice at 1 and 3 months of age. Representative full-cornea montages are shown for each age, genotype and sex. (B) Quantification of epithelial nerve fibre volume is presented for each group (n = 5 corneas per age per sex per genotype). Individual data points and statistical comparisons between genotypes are shown. (C) Corneal sensitivity measurements in female and male Pitx2⁺/⁺ and Pitx2^egl1/egl1^ mice at 1 and 3 months of age. Bar graphs display mean values with individual data points for each cohort (n = 5 and n = 10 per sex per genotype at 1 and 3 months respectively). Statistical significance between genotypes is indicated as ns (not significant), *p < 0.05, **p < 0.01, ***p < 0.001, ****p < 0.0001.

Consistent with these structural changes, corneal sensitivity revealed sex-dependent differences. At 3-month, mutant males (3.72 g^-1^) had reduced responses compared to control males (8.09 g^-1^p = 0.021), while females remained comparable across genotypes (Fig 2C). The disappearance of normal sex differences in the mutant group indicates that the Pitx2 gene mutation affects corneal sensory function differently depending on sex. Given the emergence of structural and functional alterations at this stage, we then sought to investigate corneal transcriptomic changes at 3 months of age to identify the molecular pathways associated with these deficits. Because the phenotype became evident at 3 months, we then focused our molecular analyses on this stage to determine whether sex-specific transcriptional programs in the cornea could account for the structural and functional deficits observed.

### 3.3 Baseline corneal transcription shows limited sex-dependent differences

To place the mutant phenotypes in context, we first defined the extent of baseline sexual dimorphism in the healthy cornea. Transcriptomic analysis of control mice revealed only modest significant differences between females and males, with 62 genes differentially expressed (Fig. 3A). The most enriched biological processes (Fig. 3B) involved chromatin organization and histone modification (H3-K4 and H3-K27 demethylation), translational regulation (such as positive regulation of translational fidelity and formation of the translation preinitiation complex), and endoplasmic reticulum stress (ER) responses (including IRE1-mediated unfolded protein response and ER-associated degradation). KEGG pathway analysis showed no significant enrichment for most pathways; however, protein processing in the ER was enriched among upregulated genes (Fig. S3A–B).

**Figure 3:**
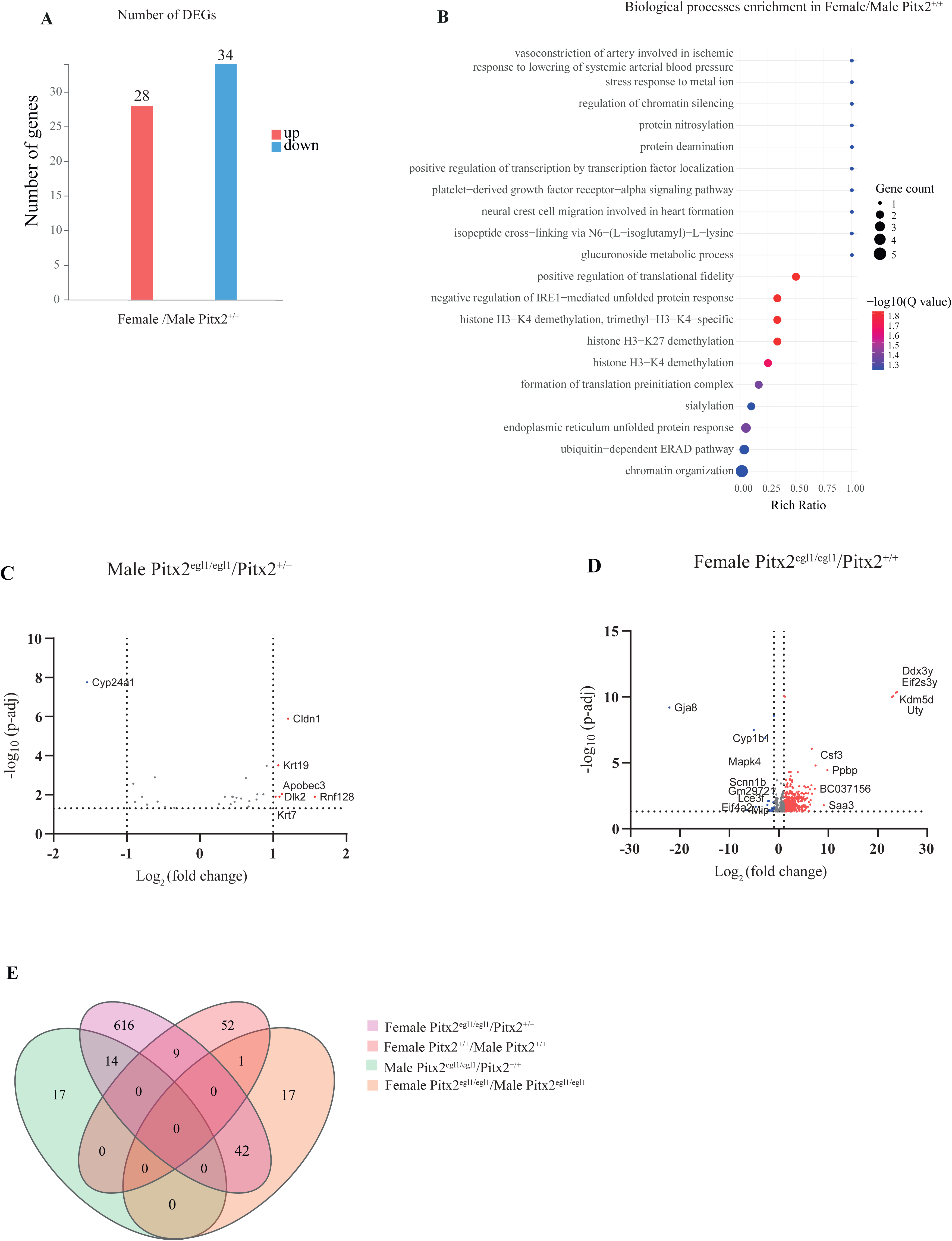
Differential gene expression and functional enrichment analysis in female versus male Pitx2*^+/+^* and in male and female Pitx2^egl1/egl1^ versus Pitx2*^+/+^* corneas. (A) Differentially expressed genes (DEGs) identified between female and male Pitx2⁺/⁺ corneas. Upregulated genes in females are shown in red, and downregulated genes are shown in blue. A total of 28 genes were upregulated and 34 genes were downregulated. (B) Gene Ontology (GO) biological process enrichment analysis of DEGs. The y-axis represents enriched GO biological process terms, and the x-axis represents the rich ratio. Bubble size indicates gene count (1–5 genes per term), and bubble color reflects statistical significance, expressed as - log10(Q value), ranging from blue (less significant) to red (more significant). (C) Volcano plot showing differentially expressed genes (DEGs) in male Pitx2^egl1/egl1^ versus male Pitx2⁺/⁺ corneas. The x-axis represents log2 fold change, and the y-axis represents −log10 (adjusted p-value). Vertical dashed lines indicate log2 fold change thresholds of −1 and 1, and the horizontal dashed line indicates a significance threshold of −log10 (p-adjusted) = 1.3. Genes with log2 fold change < −1 are shown in blue (downregulated), and genes with log2 fold change > 1 are shown in red (upregulated). (D) Volcano plot showing DEGs in female Pitx2^egl1/egl1^ versus female Pitx2⁺/⁺ mice, displayed as in (C). (E) Venn diagram illustrating overlaps of differentially expressed genes between comparisons: male Pitx2^egl1/egl1^ versus male Pitx2⁺/⁺, female Pitx2^egl1/egl1^ versus female Pitx2⁺/⁺, female Pitx2⁺/⁺ versus male Pitx2⁺/⁺, and female Pitx2^egl1/egl1^ versus male Pitx2^egl1/egl1^.

Overall, these results indicate that baseline transcriptional differences between male and female corneas are limited. Complete DEG and enrichment datasets are provided in Supplementary Tables S1–S2.

### 3.4 Pitx2 mutation amplifies sex-specific transcriptional differences in the cornea

We next asked how Pitx2 mutation reshapes the corneal transcriptome in each sex. To address this, we compared mutant and control corneas separately in males and females. Volcano plots (Fig. 3C–D) immediately revealed a marked asymmetry in the magnitude of the response, and Venn diagram analysis confirmed minimal overlap between the resulting DEG sets (Fig. 3E), indicating that the transcriptional consequences of the mutation are strongly sex dependent. Thus, whereas baseline sex differences in the cornea were modest, Pitx2 mutation markedly widened the transcriptional gap between females and males, with females showing a broader and largely distinct response. To understand the nature of this divergence, we then analysed the female and male mutant corneas separately.

#### 3.4.1 Pitx2 mutation in female cornea induces extensive differential gene expression and activation of inflammatory signaling pathways

Consistent with the global comparison above, female corneas exhibited the most pronounced transcriptional response to Pitx2 deficiency. In females, the mutation triggered extensive transcriptional reprogramming, with 681 DEGs in total, including 593 upregulated and 88 downregulated genes (Fig. 4A), indicating a response dominated by gene activation. The top 15 up- and downregulated genes are listed in Table 1. Significant enrichment was observed in biological processes related to cell migration, proliferation, apoptosis, and tissue remodeling. Immune-related and inflammatory pathways, including inflammatory response (Q = 5.874658E^-5^), innate immune response (Q = 4.351522E^-4^), and cellular response to interleukin-1 (Q = 5.270612E^-4^), were also prominently upregulated (Fig. 4C), with several top upregulated genes such as *Ppbp* (∼907-fold, Q = 3.63 × 10⁻^5^)*, Saa3* (∼545-fold, Q = 0.0167)*, Ecm1* (∼81-fold, Q = 0.0150), and *Ltf* (∼79-fold, Q = 0.00104). mapping to these processes. In contrast, only a limited number of downregulated genes, including *Mip, Cyp1b1, Scnn1b,* and *Siglecg*, were associated with enriched biological processes. KEGG pathway analysis highlighted significant upregulated pathways, including IL-17, TNF, NF-κB, and MAPK signaling, indicating activation of pro-inflammatory and stress-response networks (Fig. 5A). Downregulated pathways were not statistically significant (Fig. 5B). Altogether, these data show that female mutant corneas mount a broad transcriptional response centred on inflammation, stress signalling, tissue remodelling, and epithelial activation. Complete DEG and enrichment datasets are provided in Supplementary Tables S3–S4. We next investigated whether male corneas showed a comparable response or instead engaged a distinct molecular program.

**Figure 4:**
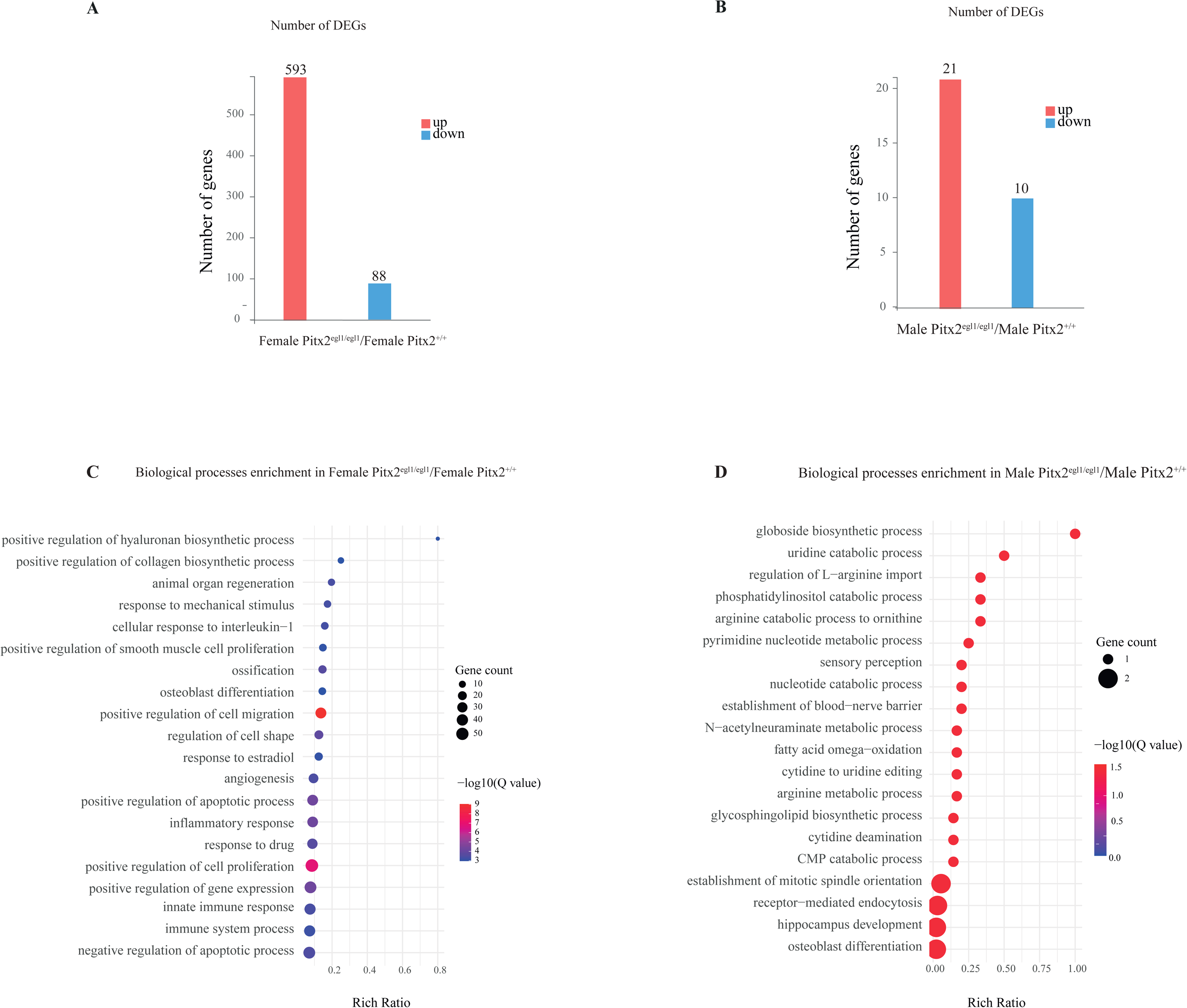
Differential gene expression and functional enrichment analysis in female and male Pitx2^egl1/egl1^ versus Pitx2*^+/+^* corneas. (A) Differentially expressed genes (DEGs) identified between female Pitx2^egl1/egl1^ and Pitx2^+/+^ corneas. Upregulated genes in Pitx2^egl1/egl1^ are shown in red, and downregulated genes are shown in blue. A total of 593 genes were upregulated and 88 genes were downregulated. (B) Differentially expressed genes (DEGs) identified between male Pitx2^egl1/egl1^ and Pitx2^+/+^ corneas. Upregulated genes in Pitx2^egl1/egl1^ are shown in red, and downregulated genes are shown in blue. A total of 21 genes were upregulated and 10 genes were downregulated. (C) Gene Ontology (GO) biological process enrichment analysis of DEGs. The y-axis represents enriched GO biological process terms, and the x-axis represents the rich ratio. Bubble size indicates gene count (10–50 genes per term), and bubble color reflects statistical significance, expressed as - log10(Q value), ranging from blue (less significant) to red (more significant). (D) Gene Ontology (GO) biological process enrichment analysis of DEGs. The y-axis represents enriched GO biological process terms, and the x-axis represents the rich ratio. Bubble size indicates gene count (1 or 2 genes per term), and bubble color reflects statistical significance, expressed as - log10(Q value), ranging from blue (less significant) to red (more significant).

**Figure 5:**
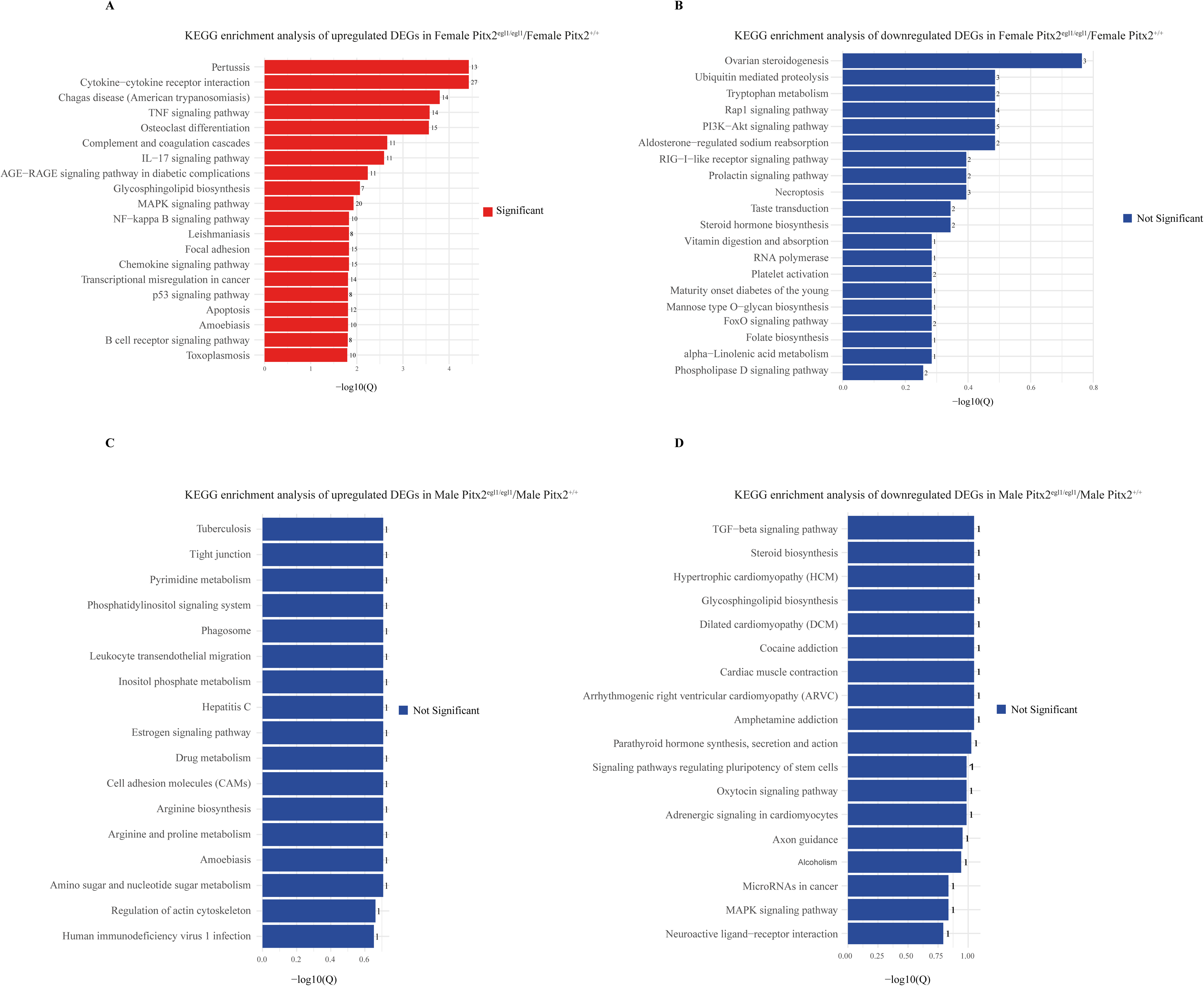
Functional enrichment analysis (KEGG pathway) in female and male Pitx2^egl1/egl1^ versus Pitx2*^+/+^* corneas. (A-B) KEGG pathway enrichment analysis of upregulated and downregulated DEGs respectively in female Pitx2^egl1/egl1^ versus Pitx2 ^+/+^. (C-D) KEGG pathway enrichment analysis of upregulated and downregulated DEGs respectively in male Pitx2^egl1/egl1^ versus Pitx2 ^+/+^. Red bars indicate significantly enriched pathways, while blue bars indicate non-significant pathways. The number of DEGs associated with each pathway is indicated.

#### 3.4.2 Male corneas show restricted gene expression changes in response to Pitx2 mutation

In contrast to females, male corneas displayed a much more limited transcriptional response to Pitx2 mutation. Only 31 DEGs were identified in males, including 21 upregulated and 10 downregulated genes (Fig. 4B), and the top 15 up- and downregulated genes are presented in Table 2. Rather than broad inflammatory activation, enriched biological processes in males were mainly related to metabolic regulation and selected sensory-associated functions, including nucleotide and amino acid metabolism (*Upp1*: uridine catabolic, CMP catabolic, and pyrimidine nucleotide metabolic processes; *Arg1*: arginine metabolic processes), glycosphingolipid and globoside biosynthesis (*A4galt* downregulated), sensory perception (*Pdyn* downregulated), and establishment of the blood–nerve barrier (*Cldn1* upregulated, FC = 2.3). Additional metabolic adjustments included downregulation of Vitamin D metabolism (*Cyp24a1* downregulated) and cytidine deamination/editing pathways (*Apobec3* upregulated), highlighting discrete transcriptional modulation rather than broad tissue remodeling (Fig. 4D). KEGG pathways in males were not significant (Fig. 5C–D). Supplementary data (Supplementary Tables S5-S6) summarize all DEGs in male corneas and the results of biological process enrichment analyses.

Direct comparison of female and male mutants revealed a sex-specific transcriptional divergence. In females, there was a robust activation of stress-response and immune pathways, including genes linked to inflammatory signaling and neutrophil aggregation and siderophore transport (Fig. S4A). KEGG pathway analysis highlighted a significant upregulation of the IL-17 signaling pathway (Q < 0.00631), supporting a strong pro-inflammatory response (Fig. S4B). Concurrently, several metabolic and homeostatic pathways were comparatively downregulated in females relative to males, including purine and pyrimidine metabolism, nicotinamide riboside catabolic processes, and acyl-CoA metabolism. Sex-specific differences were also observed in cell cycle regulation and DNA integrity, with female-specific enrichment of cell division processes and histone H2A-S139 phosphorylation (Fig. S4C).

We therefore next examined the trigeminal ganglia to determine whether these sex-specific corneal phenotypes were accompanied by corresponding changes in upstream sensory neurons.

### 3.5 Trigeminal ganglia in Pitx2⁺/⁺ mice show only limited sex differences in cellular and neuronal organization

Transcriptomic analysis of control trigeminal ganglia revealed only subtle sex-dependent differences, with 30 genes upregulated and 21 downregulated in males relative to females (Fig. 6A). Enrichment analysis pointed mainly to modest variation in translational machinery and intracellular neuronal organization, including components of eukaryotic initiation factor complexes and axonal or somatodendritic compartments (Fig. 6B). No KEGG pathways reached statistical significance (Fig. S5A-B). Complete lists of DEGs and enrichment results can be found in Supplementary Table S7-S8.

**Figure 6:**
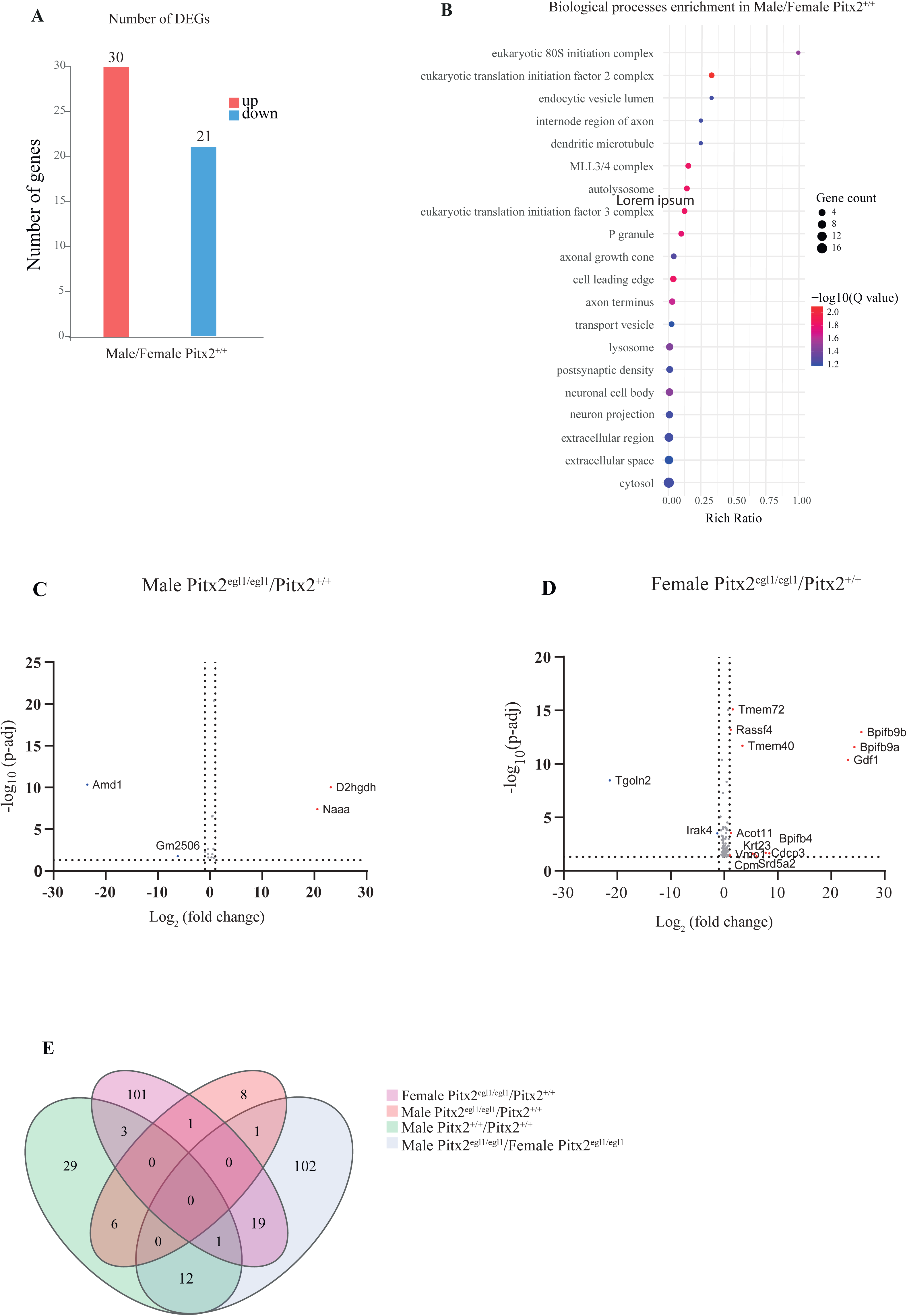
Differential gene expression and functional enrichment analysis in male versus female Pitx2*^+/+^* and in male and female Pitx2^egl1/egl1^ versus Pitx2^+/+^ trigeminal ganglia. (A) Differentially expressed genes (DEGs) identified between female and male Pitx2⁺/⁺ trigeminal ganglia. Upregulated genes in females are shown in red, and downregulated genes are shown in blue. A total of 30 genes were upregulated and 21 genes were downregulated. (B) Gene Ontology (GO) biological process enrichment analysis of DEGs. The y-axis represents enriched GO biological process terms, and the x-axis represents the rich ratio. Bubble size indicates gene count (4-16 genes per term), and bubble color reflects statistical significance, expressed as - log10(Q value), ranging from blue (less significant) to red (more significant). (C) Volcano plot showing differentially expressed genes (DEGs) in male Pitx2^egl1/egl1^ versus male Pitx2⁺/⁺ trigeminal ganglia. The x-axis represents log2 fold change, and the y-axis represents −log10 (adjusted p-value). Vertical dashed lines indicate log2 fold change thresholds of −1 and 1, and the horizontal dashed line indicates a significance threshold of −log10 (p-adjusted) = 1.3. Genes with log2 fold change < −1 are shown in blue (downregulated), and genes with log2 fold change > 1 are shown in red (upregulated). (D) Volcano plot showing DEGs in female Pitx2^egl1/egl1^ versus female Pitx2⁺^/^⁺ mice, displayed as in (C). (E) Venn diagram illustrating overlaps of differentially expressed genes between comparisons: male Pitx2^egl1/egl1^ versus male Pitx2⁺^/^⁺, female Pitx2^egl1/egl1^ versus female Pitx2⁺^/^⁺, male Pitx2⁺/⁺ versus female Pitx2⁺/⁺, and male Pitx2^egl1/egl1^ versus female Pitx2^egl1/egl1^.

These results indicate that, as in the cornea, baseline sexual dimorphism in the trigeminal ganglia is limited and does not involve major pathway-level reprogramming.

We next examined whether Pitx2 mutation indirectly affects trigeminal ganglia in a sex-dependent manner. Although Pitx2 is expressed during craniofacial development and in neural crest-derived periocular tissues, there is no clear evidence for its expression in differentiated adult trigeminal ganglia neurons, suggesting that any transcriptomic changes observed here are more likely secondary to altered peripheral target tissues than cell-autonomous effects in the ganglion itself [5,32,33]. Supporting this idea, Figure S6 shows sex-dependent differences between male and female Pitx2^egl1/egl1^ mice with differential enrichment of ribosome and translation-associated biological processes, as well as variation in prolactin signaling pathways.

Separate analysis of male and female Pitx2^egl1/egl1^ and Pitx2^+/+^ trigeminal ganglia by volcano plots (Fig. 6C–D) and overlap analysis revealed clear sex-related differences in the number and identity of DEGs (Fig. 6E). We therefore analysed female and male trigeminal ganglia separately to define the biological processes and pathways associated with these distinct responses.

### 3.6 Pitx2 mutation leads to coordinated adaptations of neurons and axons in the female trigeminal ganglia associated with corneal innervation

In keeping with the relatively restricted corneal phenotype in females, female trigeminal ganglia underwent substantially broader transcriptional reprogramming in response to Pitx2 mutation. In female trigeminal ganglia, 125 DEGs were identified, including 68 upregulated and 57 downregulated genes (Fig. 7A), and the top 20 up- and downregulated genes are listed in Table 3. Upregulated genes such as *Snap25, Prkaa2, Hpca, Atg7, Srd5a2*, and *Acot11* were enriched in axonal compartments, cytoskeletal organisation, and neuronal cell body-associated terms (Fig. 7C; Table S13), pointing to changes in neuronal architecture and excitability. In parallel, C-fiber–associated channels (Scn10a, Scn11a), ribosomal components (*Rps6, Rpl5, Rpl38, Rps3a1*) were relatively reduced, consistent with KEGG pathway analysis showing downregulation of the ribosome pathway significantly, suggesting a shift in translational dynamics (Fig. 8B), whereas no significant pathways were identified for the upregulated genes (Fig. 8A). Downregulated genes such as *Anxa4, Rlbp1, Osbpl10, Prkca,* and *Baiap2l1* further pointed to alterations in cytoskeletal organization, membrane trafficking, and signal transduction (Fig.7C, Table S9).

**Figure 7:**
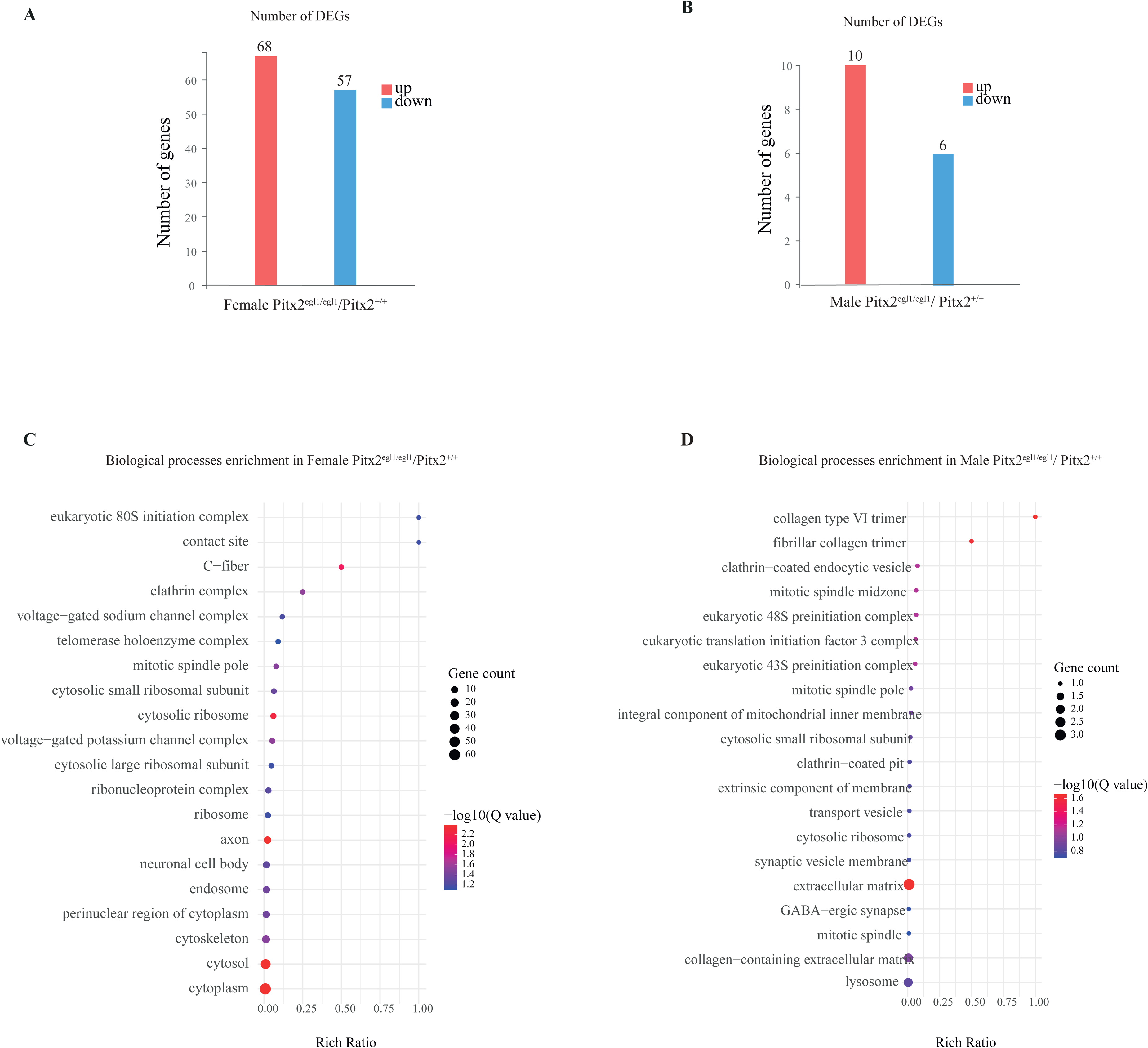
Differential gene expression and functional enrichment analysis in female Pitx2^egl1/egl1^ versus Pitx2^+/+^ trigeminal ganglia. (A) Differentially expressed genes (DEGs) identified between female Pitx2^egl1/egl1^ and Pitx2^+/+^ trigeminal ganglia. Upregulated genes in Pitx2egl1/egl1 are shown in red, and downregulated genes are shown in blue. A total of 68 genes were upregulated and 57 genes were downregulated. (B) Differentially expressed genes (DEGs) identified between male Pitx2^egl1/egl1^ and Pitx2^+/+^ trigeminal ganglia. Upregulated genes in Pitx2^egl1/egl1^ are shown in red, and downregulated genes are shown in blue. A total of 10 genes were upregulated and 6 genes were downregulated. (C) Gene Ontology (GO) biological process enrichment analysis of DEGs. The y-axis represents enriched GO biological process terms, and the x-axis represents the rich ratio. Bubble size indicates gene count (10–60 genes per term), and bubble color reflects statistical significance, expressed as - log10(Q value), ranging from blue (less significant) to red (more significant). (D) Gene Ontology (GO) biological process enrichment analysis of DEGs. The y-axis represents enriched GO biological process terms, and the x-axis represents the rich ratio. Bubble size indicates gene count (1-3 genes per term), and bubble color reflects statistical significance, expressed as - log10(Q value), ranging from blue (less significant) to red (more significant).

**Figure 8:**
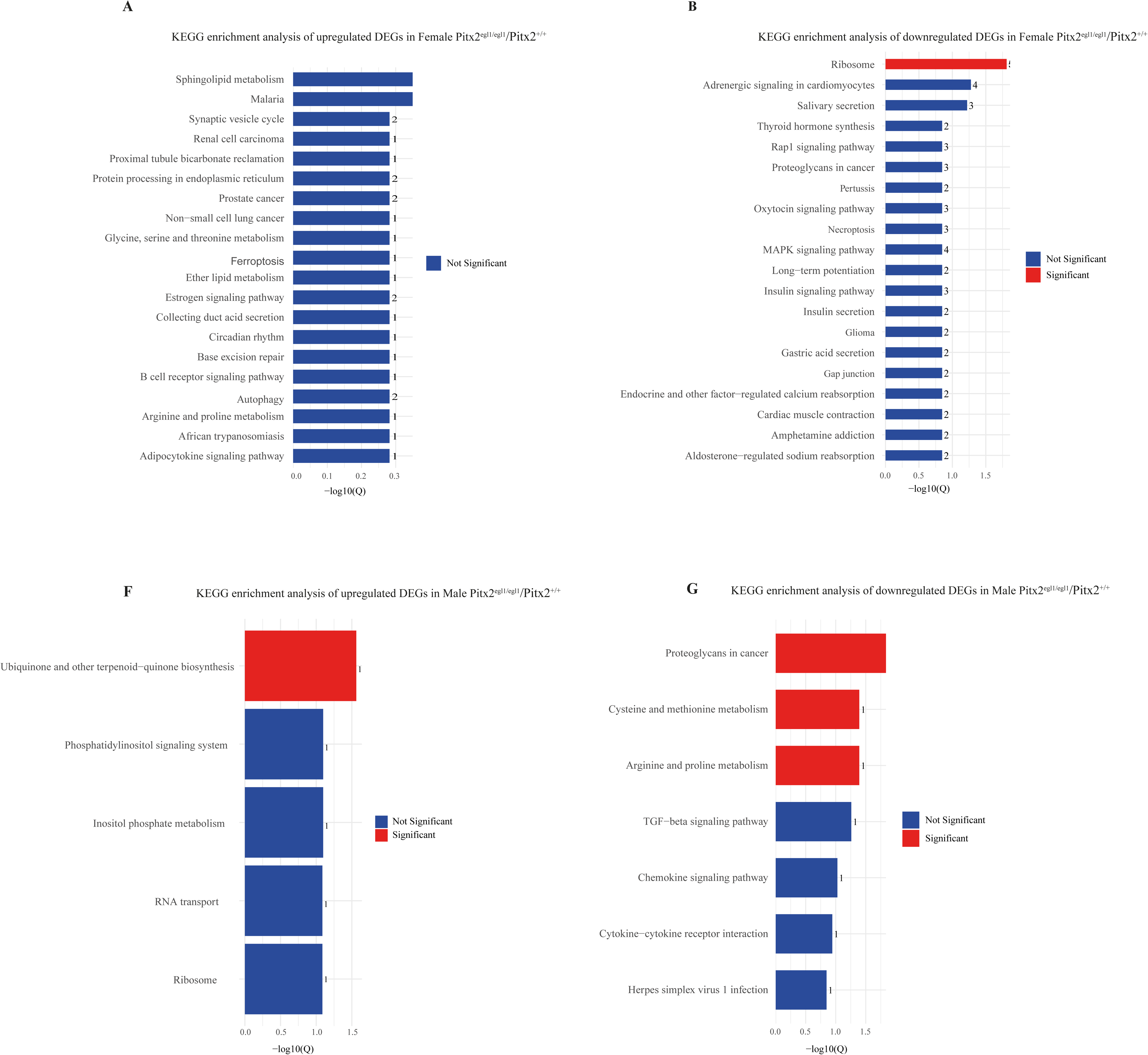
Functional enrichment analysis (KEGG pathway) in female and male Pitx2^egl1/egl1^ versus Pitx2*^+/+^* trigeminal ganglia. (A-B) KEGG pathway enrichment analysis of upregulated and downregulated DEGs respectively in female Pitx2^egl1/egl1^ versus Pitx2 ^+/+^. (C-D) KEGG pathway enrichment analysis of upregulated and downregulated DEGs respectively in male Pitx2^egl1/egl1^ versus Pitx2 ^+/+^. Red bars indicate significantly enriched pathways, while blue bars indicate non-significant pathways. The number of DEGs associated with each pathway is indicated.

We next investigated whether male trigeminal ganglia exhibited a similarly extensive adaptation or a modest response.

### 3.7 Male trigeminal ganglia present limited upstream transcriptional adjustments supporting corneal structure and metabolism

By contrast, with the relatively restricted corneal phenotype in males, male trigeminal ganglia showed only modest transcriptional changes in response to Pitx2 mutation. We identified 16 DEGs in male trigeminal ganglia, including 10 upregulated and 6 downregulated genes (Fig. 7B; Table 4). GO enrichment analysis highlighted extracellular matrix-related components (Q value = 0.02; rich ratio = 0.01) (Fig. 7D; Table S10), suggesting subtle changes in tissue support and structural organisation. It should be noted that these ECM-related processes mirror those observed in the cornea, including structural remodeling, glycosphingolipid metabolism, and vesicle trafficking, suggesting coordinated regulation of tissue structure along the cornea–sensory neuron axis. Downregulated genes, such as *Amd1* (∼907-fold, Q = 8,3E^-8^), indicate reduced amino acid metabolism, reflected in KEGG pathways for cysteine/methionine and arginine/proline metabolism (Fig. 8F-G), which also correspond to corneal metabolic processes. Overall, the male trigeminal ganglia response was limited and was primarily characterised by modest changes in extracellular matrix and amino acid metabolic pathways. These alterations parallel the restrained corneal transcriptional response observed in males and are consistent with a mild upstream adaptation that may help shape corneal structural and metabolic homeostasis without triggering broad neuronal reprogramming.

Taken together, these results show that female trigeminal ganglia respond to Pitx2-associated glaucoma through broader neuronal and axonal reprogramming than males, whereas male TG exhibit only limited metabolic and structural adjustments.

## 4. Discussion

In this study, we show that the Pitx2^egl1/egl1^ mouse reproduces the major features of early-onset glaucoma and, in addition, develops progressive abnormalities of the corneal sensory system. Ocular hypertension, fundus alterations, reduced visual evoked potential amplitudes, and optic nerve degeneration confirmed the relevance of the model for developmental glaucoma. Against this background, our results identify a corneal phenotype that has received little attention in glaucoma research: a progressive reduction in corneal epithelial nerve fiber volume associated with sex-dependent molecular responses in both the cornea and trigeminal ganglia. Importantly, corneal nerve loss was not detected at 1 month but became evident at 3 months, indicating that it emerges during disease progression rather than reflecting only an early developmental defect.

The main novelty of the study lies in the demonstration that glaucoma in this model is associated with corneal denervation. This is the first study to specifically examine the impact of developmental glaucoma on corneal innervation by combining ocular phenotyping, corneal transcriptomics, trigeminal ganglion transcriptomics, structural nerve analysis, and sensory testing. This is important as glaucoma is usually considered primarily through the lens of retinal ganglion cell loss and optic nerve damage [34,35]. Our data suggest that the disease may also affect the corneal sensory apparatus, a system that is essential for epithelial maintenance, wound healing, and protective ocular surface reflexes [8,9,11,36]. Although direct extrapolation to patients must remain cautious, this observation raises the possibility that corneal sensory integrity may represent an underappreciated component of disease burden in developmental glaucoma.

A second key finding is that the structural and functional readouts were not fully concordant. Both female and male mutant mice exhibited reduced corneal nerve fiber volume at 3 months, but reduced mechanical sensitivity was detected only in males. This divergence suggests that structural nerve loss does not necessarily translate into the same functional outcome in both sexes. Our data do not establish the mechanism underlying this difference, but they do indicate that females and males mount distinct tissue responses to the same disease context. At minimum, these findings argue against a uniform relationship between corneal nerve density and mechanically evoked blink responses in this model.

The transcriptomic analyses support this interpretation. In control animals, baseline sex differences were limited in both cornea and trigeminal ganglia, indicating that the marked divergence observed in mutants is unlikely to simply reflect pre-existing physiological dimorphism. In the cornea, female mutants showed broad transcriptional changes enriched in inflammatory, stress-response, and tissue-remodeling pathways, whereas male mutants showed a narrower response centered on metabolic and homeostatic processes. The trigeminal ganglion displayed a similar pattern, with broader neuronal and axonal pathway changes in females and more modest extracellular matrix and metabolic changes in males. These data do not demonstrate causality, but they support the idea that the cornea and its sensory ganglion engage coordinated yet sex-dependent responses during glaucoma progression.

The trigeminal findings are particularly informative because they suggest that corneal involvement is not an isolated local event. Corneal sensory fibers originate from trigeminal neurons, and the cornea and trigeminal ganglion form a tightly coupled biological unit [37–39]. The observation that both compartments are transcriptionally altered in mutant animals is therefore consistent with a broader disturbance of the corneal sensory axis. At the same time, the present data do not establish whether ganglionic changes are upstream drivers, downstream consequences, or parallel adaptations to chronic ocular pathology. Given the lack of clear evidence for Pitx2 expression in differentiated adult trigeminal neurons [32,40,40–42], it is reasonable to interpret these ganglionic changes cautiously as indirect responses to disease rather than direct cell-autonomous effects of the mutation.

These results may also have practical implications. Corneal nerves are central to ocular surface homeostasis, and their impairment can influence epithelial stability, regeneration, tear reflexes, and discomfort [10,36,43,44]. If similar alterations occur in patients with developmental glaucoma, corneal sensory changes could contribute to clinical manifestations that are not captured by conventional glaucoma endpoints focused on IOP and optic nerve status. Our study does not justify immediate clinical extrapolation, but it does suggest that ocular surface status, corneal sensitivity, and possibly corneal nerve integrity deserve greater attention in the broader evaluation of disease impact. In that sense, the work opens a new angle on glaucoma pathophysiology by suggesting that the corneal sensory system may be another tissue compartment affected by the disease.

This point is especially relevant because the translational value of the finding lies less in proving a mechanism than in redefining the scope of what should be monitored. In clinical practice, patients with developmental glaucoma are primarily followed for pressure control, optic nerve damage, and visual function [45–47]. Our data suggest that this framework may be incomplete. A better appreciation of corneal involvement could ultimately help refine patient follow-up, particularly in individuals with ocular surface symptoms, epithelial fragility, or suspected sensory dysfunction. Future human studies will be needed to determine whether corneal nerve imaging or esthesiometry could provide useful complementary information in this setting.

Some limitations should be considered when interpreting the present results. This study relies on a developmental mouse model, which provides a relevant experimental framework but may not encompass the full biological and clinical diversity of human glaucoma. In addition, corneal sensory function was evaluated using a mechanical assay, and complementary approaches exploring other sensory modalities would help further refine the functional characterization of the phenotype. Finally, while the data support coordinated changes in the cornea and trigeminal ganglion, additional studies will be needed to clarify the respective contribution and temporal relationship of ocular hypertension, anterior segment abnormalities, corneal denervation, and ganglion remodeling.

In summary, our data show that Pitx2-associated early-onset glaucoma is accompanied by progressive corneal denervation together with sex-dependent molecular changes in the cornea and trigeminal ganglia. Beyond extending the phenotype of this model, these findings identify the corneal sensory system as a previously overlooked target of glaucomatous pathology. This conceptual advance is important because it broadens the view of glaucoma from a disease centered on the posterior segment and optic nerve to one that may also affect anterior sensory ocular tissues, with potential implications for disease monitoring and future patient care.

## Acknowledgements

The authors acknowledge the RNA-sequencing services provided by the BGI Group unit in Hong Kong. The authors thank the MRI-DBS imaging facility, a member of the France-BioImaging national research infrastructure, supported by the French National Research Agency (ANR-10-INBS-04, “Investments for the Future”).

The authors also thank the personnel of the INM animal core facility, a member of the Réseau des Animaleries de Montpellier (RAM), for their technical support.

## Author contributions: CRediT

**Conceptualization: S.S., F.M.**

**Methodology:** S.S., L.D., M.G., M.S., V.D., C.C., C.D., B.J., F.M.

**Validation:** S.S., C.C., C.D., B.J., F.M.

**Formal Analysis:** S.S., L.D., F.M.

**Investigation:** S.S., L.D., F.M.

**Writing – Original Draft Preparation: S.S.**

**Writing – Review & Editing:** S.S., L.D., M.G., M.S., V.D., C.C., C.D., B.J., F.M.

**Supervision:** C.C., C.D., B.J., F.M.

**Project Administration:** F.M.

**Funding Acquisition:** F.M.

## Disclosure

The authors have no conflict of interest.

## Funding

This work was supported by the Association Nationale de la Recherche et de la Technologie (ANRT) under a CIFRE contract, in collaboration with Laboratoires Théa.

---

**Figure S1:**
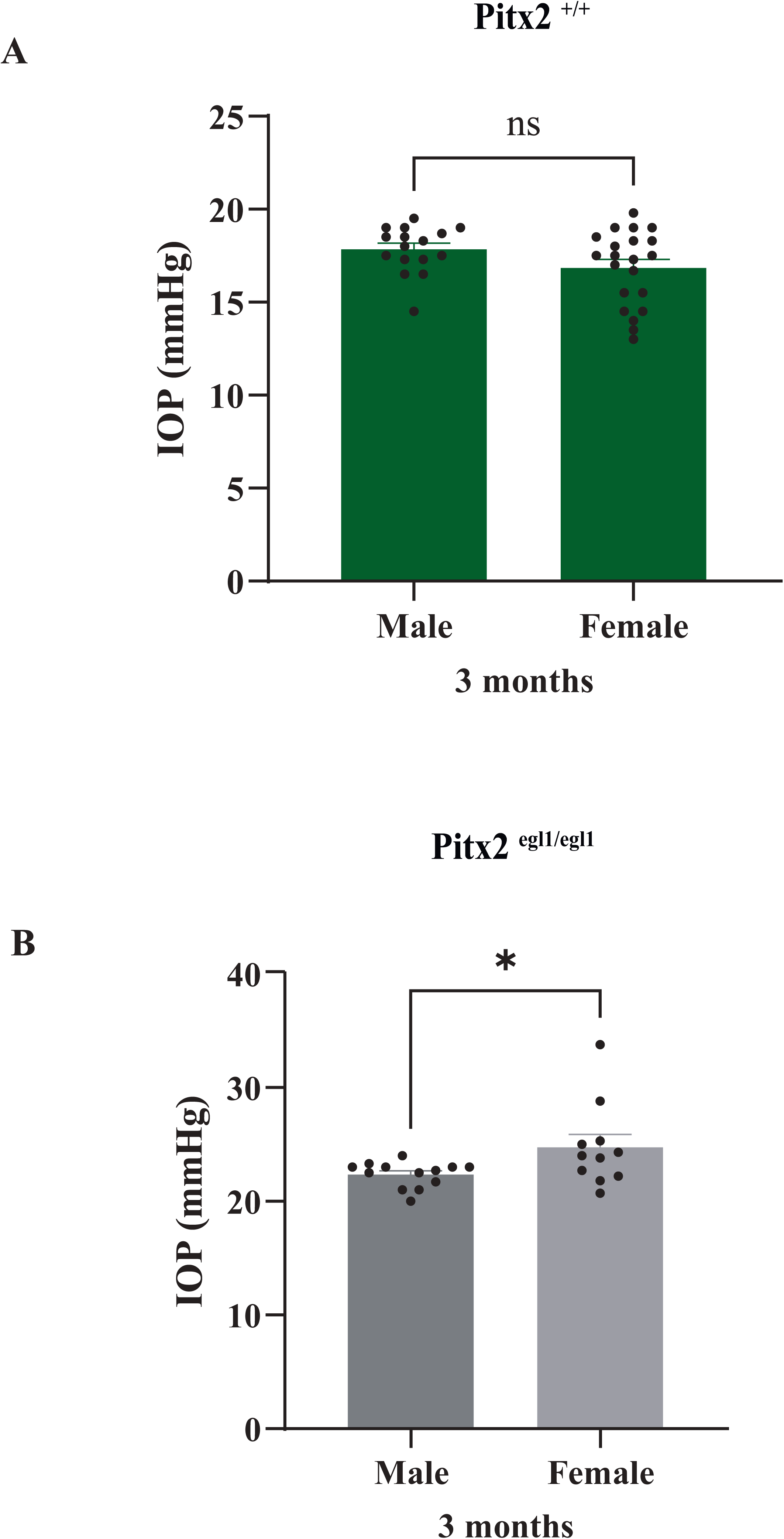
IOP measurements at 3 months in Pitx2⁺/⁺ and Pitx2^egl1/egl1^ mice. (A) IOP measurements in male and female Pitx2⁺/⁺ mice at 3 months of age. (B) IOP measurements in male and female Pitx2^egl1/egl1^ mice at 3 months of age. Pitx2⁺/⁺ (n = 37; 16 males) and Pitx2^egl1/egl1 (n = 24; 13 males). Data are presented in mmHg as mean ± SEM with individual data points shown for each group. Male and female cohorts are indicated within each genotype. Statistical significance is indicated as ns (not significant), p < 0.05, p < 0.01, *p < 0.001, and **p < 0.0001.

**Figure S2:**
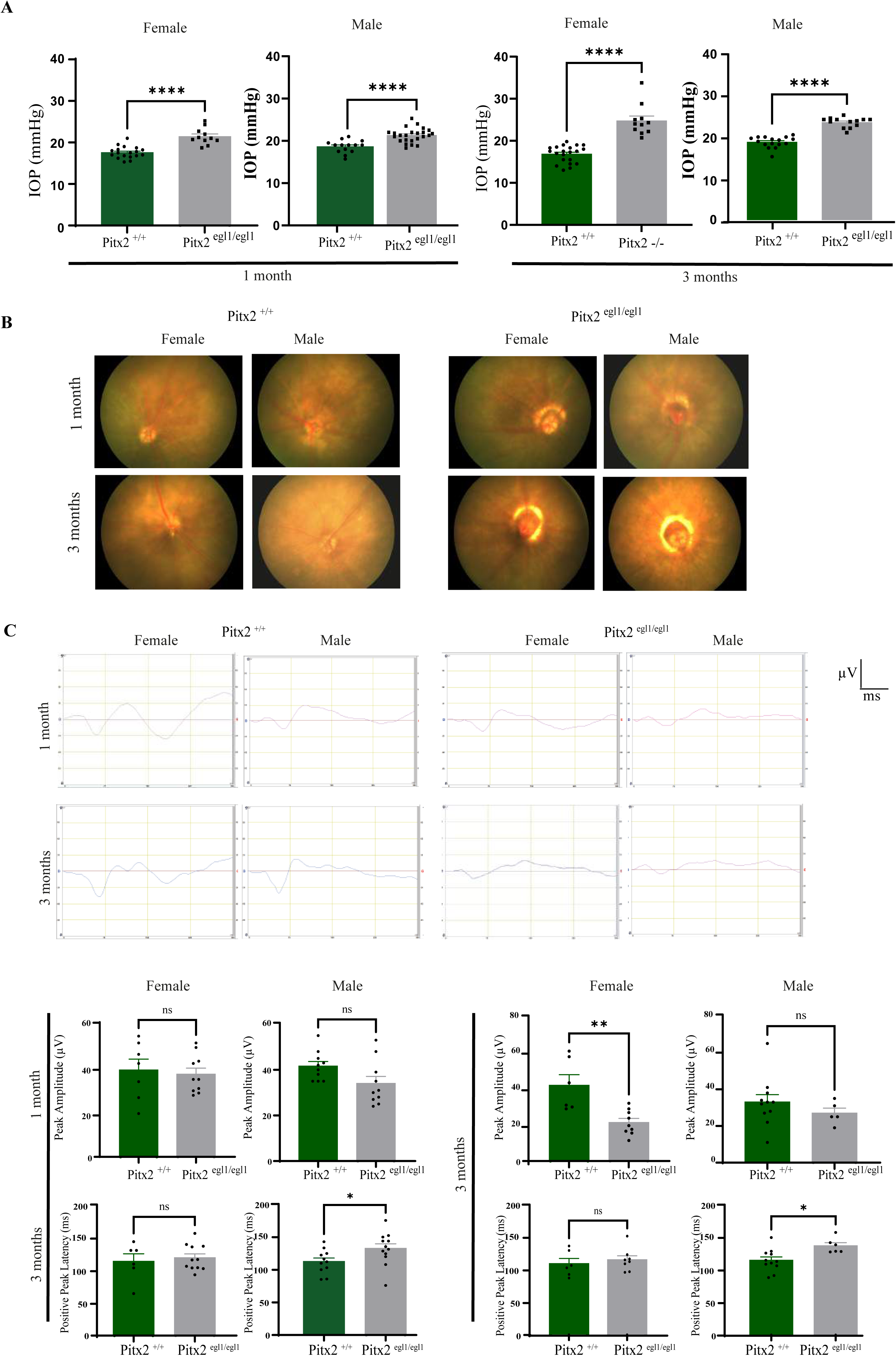
Sex-stratified ocular phenotyping and visual function in Pitx2^+/+^ and Pitx2^egl1/egl1^ mice. This figure presents the same analyses as Figure 1, with data separated by sex. (A) IOP measurements in male and female Pitx2⁺/⁺ and Pitx2^egl1/egl1^ mice at 1 and 3 months of age. Data are shown as mean ± SEM with individual data points for each group. (B) Representative fundus images from male and female Pitx2⁺/⁺ and Pitx2^egl1/egl1^ mice at 1 and 3 months of age. (C) Representative visual evoked potential (VEP) waveforms recorded in male and female Pitx2⁺/⁺ and Pitx2⁻/⁻ mice at 1 and 3 months of age, illustrating sex-specific differences in visual function. Quantification of VEP parameters, including latency and amplitude, in male and female mice across genotypes and ages. Data are presented as mean ± SEM with individual data points shown for each group. Statistical significance is indicated as ns (not significant), *p < 0.05, **p < 0.01, ***p < 0.001, ****p < 0.0001.

**Figure S3:**
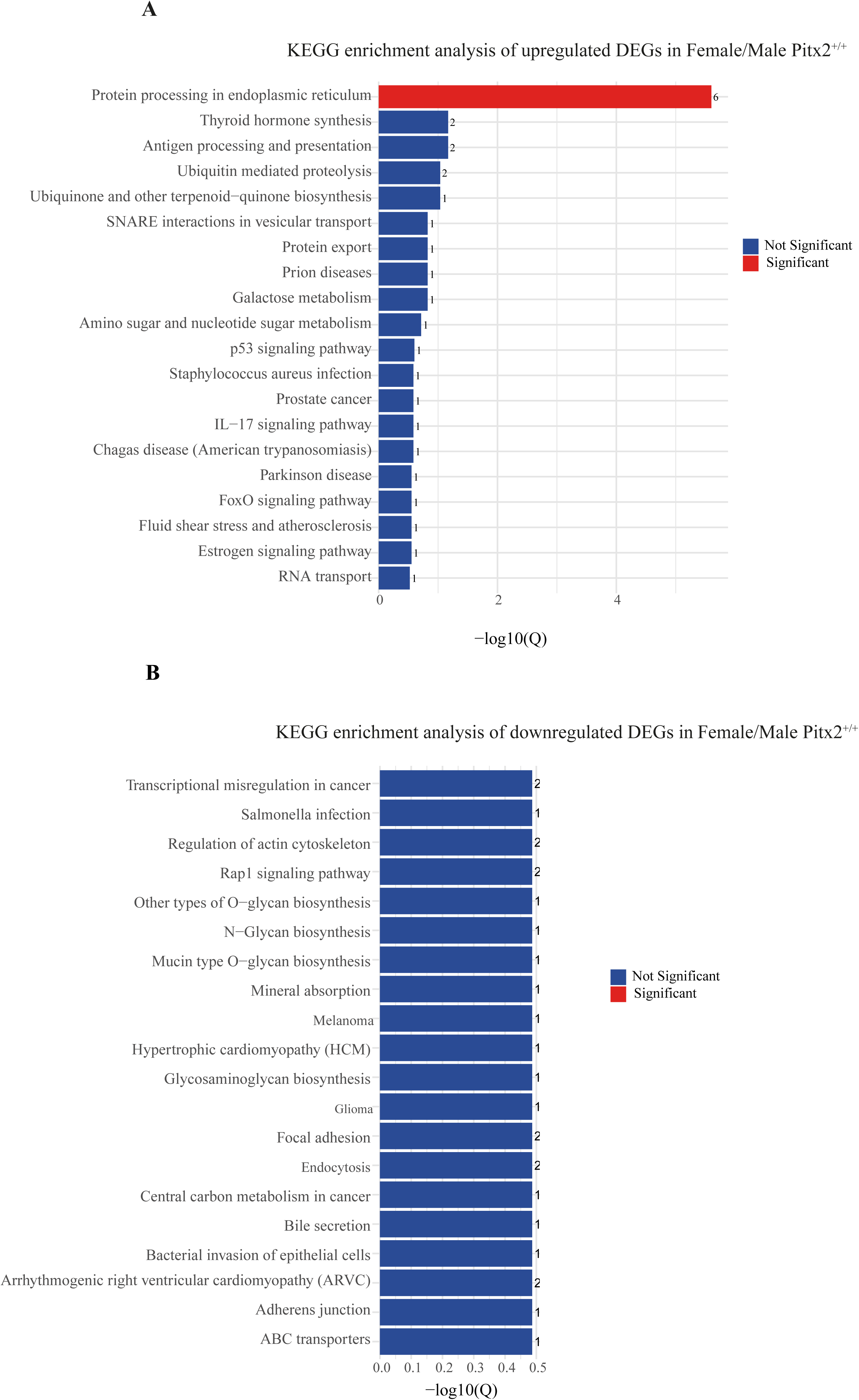
KEGG pathway analysis in female versus male Pitx2^+/+^ corneas. (A) KEGG pathway enrichment analysis of upregulated DEGs. Red bars indicate significantly enriched pathways, while blue bars indicate non-significant pathways. The number of DEGs associated with each pathway is indicated. (B) KEGG pathway enrichment analysis of downregulated DEGs, displayed as in (A).

**Figure S4:**
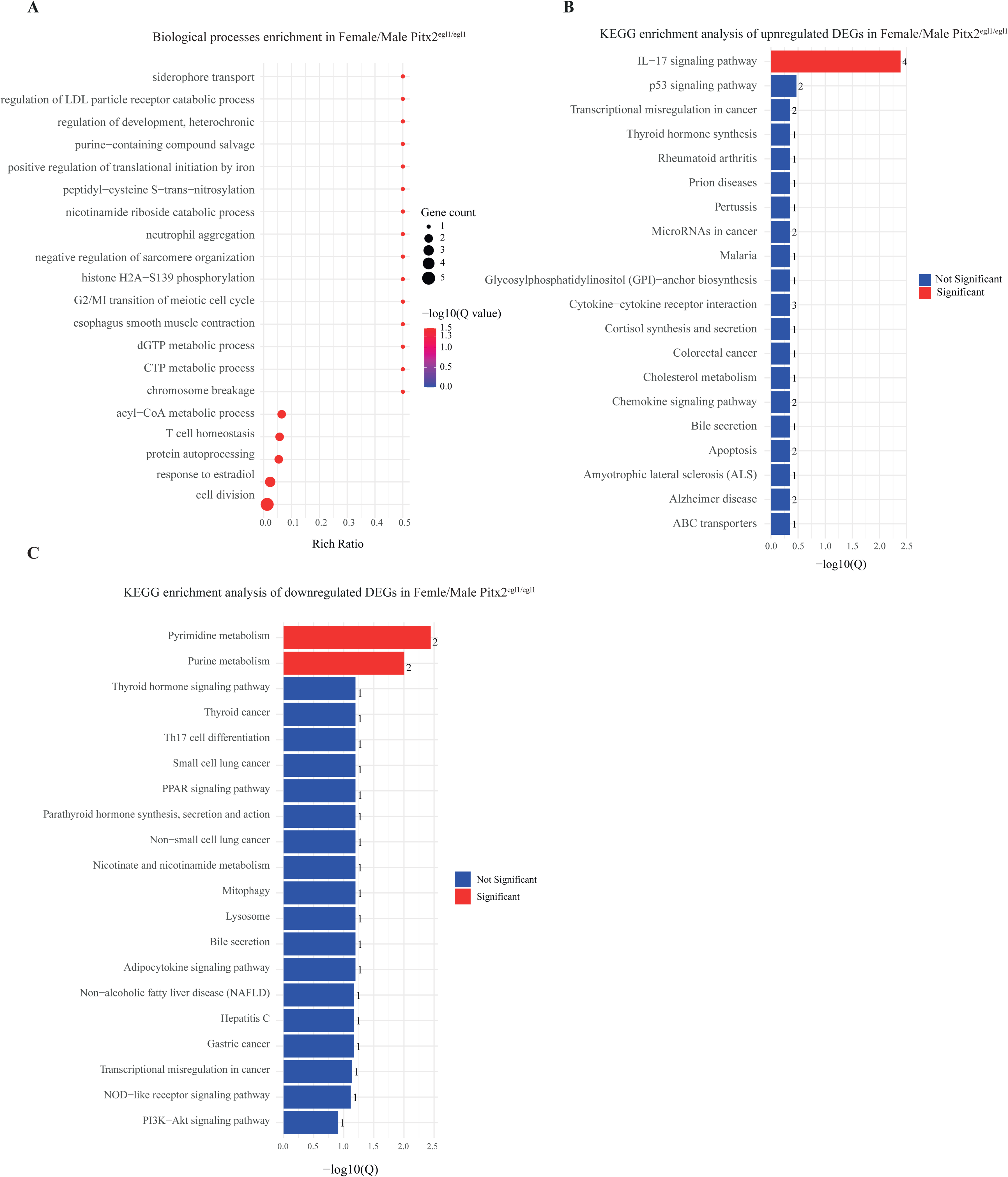
Differential gene expression and functional enrichment analysis in female versus male Pitx2^egl1/egl1^ corneas. (A) Gene Ontology (GO) biological process enrichment analysis of DEGs. The y-axis represents enriched GO biological process terms, and the x-axis represents the rich ratio. Bubble size indicates gene count (1 -5 genes per term), and bubble color reflects statistical significance, expressed as - log10(Q value), ranging from blue (less significant) to red (more significant). (B) KEGG pathway enrichment analysis of upregulated DEGs. Red bars indicate significantly enriched pathways, while blue bars indicate non-significant pathways. The number of DEGs associated with each pathway is indicated. (C) KEGG pathway enrichment analysis of downregulated DEGs, displayed as in (B).

**Figure S5:**
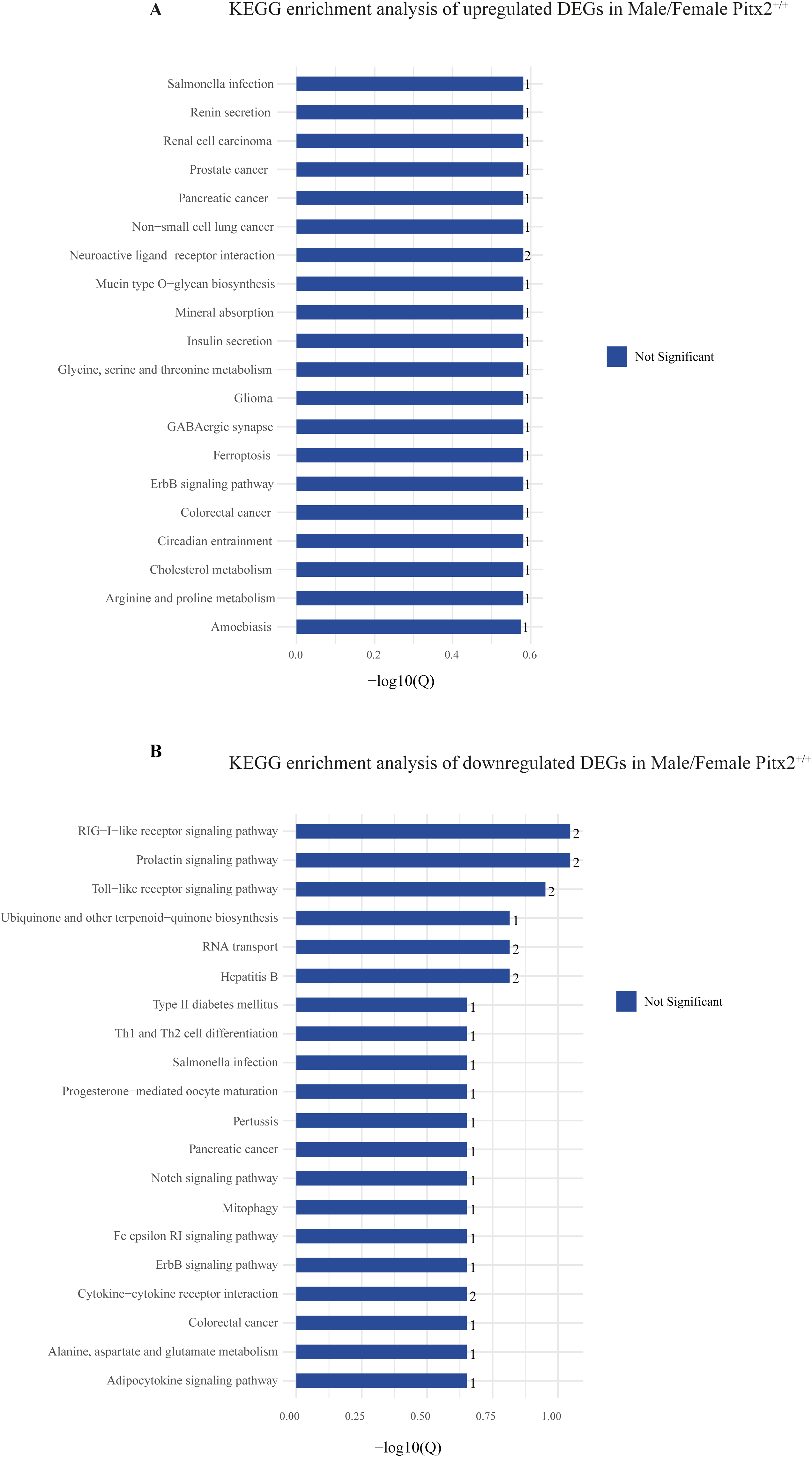
KEGG pathway analysis in male versus female Pitx2^+/+^ trigeminal ganglia. (A) KEGG pathway enrichment analysis of upregulated DEGs. Blue bars indicate non-significant pathways. The number of DEGs associated with each pathway is indicated. (B) KEGG pathway enrichment analysis of downregulated DEGs, displayed as in (A).

**Figure S6:**
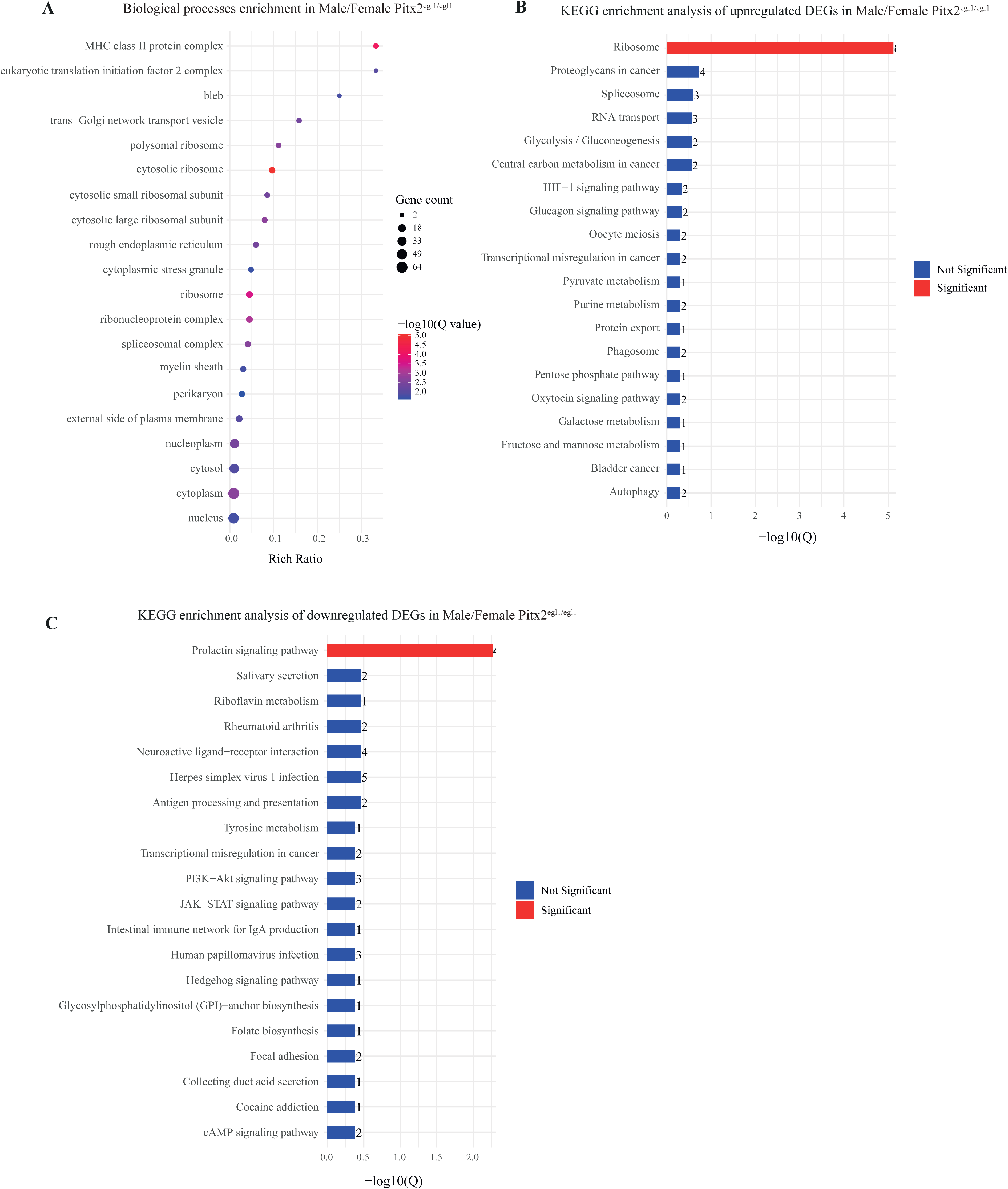
Differential gene expression and functional enrichment analysis in male versus female Pitx2^egl1/egl1^ trigeminal ganglia. (A) Gene Ontology (GO) biological process enrichment analysis of DEGs. The y-axis represents enriched GO biological process terms, and the x-axis represents the rich ratio. Bubble size indicates gene count (2-64 genes per term), and bubble color reflects statistical significance, expressed as - log10(Q value), ranging from blue (less significant) to red (more significant). (B) KEGG pathway enrichment analysis of upregulated DEGs. Red bars indicate significantly enriched pathways, while blue bars indicate non-significant pathways. The number of DEGs associated with each pathway is indicated. (C) KEGG pathway enrichment analysis of downregulated DEGs, displayed as in (B).

**Table 1: Top differentially expressed genes (DEGs) in female Pitx2^egl1/egl1^ versus Pitx2^+/+^ corneas.** This table lists the top 15 upregulated and top 15 downregulated DEGs identified in female Pitx2^egl1/egl1^ compared with Pitx2⁺/⁺ corneas. Genes were ranked based on log2 fold change. For each gene, the Gene ID, gene name, and functional description are provided, along with log2 fold change, Q value, and direction of regulation (upregulated or downregulated).

**Table 2: Top differentially expressed genes (DEGs) in male Pitx2^egl1/egl1^ versus Pitx2+/+ corneas.** This table lists the top 15 upregulated and top 15 downregulated DEGs identified in male Pitx2^egl1/egl1^ compared with Pitx2⁺/⁺ corneas. Genes were ranked based on log2 fold change. For each gene, the Gene ID, gene name, and functional description are provided, along with log2 fold change, Q value, and direction of regulation (upregulated or downregulated).

**Table 3: Top differentially expressed genes (DEGs) in female Pitx2^egl1/egl1^ versus Pitx2^+/+^ trigeminal ganglia.** This table lists the top 15 upregulated and top 15 downregulated DEGs identified in female Pitx2egl1/egl1 compared with Pitx2⁺/⁺ trigeminal ganglia. Genes were ranked based on log2 fold change. For each gene, the Gene ID, gene name, and functional description are provided, along with log2 fold change, Q value, and direction of regulation (upregulated or downregulated).

**Table 4: Top differentially expressed genes (DEGs) in male Pitx2^egl1/egl1^ versus Pitx2^+/+^ trigeminal ganglia.** This table lists the top 15 upregulated and top 15 downregulated DEGs identified in male Pitx2^egl1/egl1^ compared with Pitx2⁺/⁺ trigeminal ganglia. Genes were ranked based on log2 fold change. For each gene, the Gene ID, gene name, and functional description are provided, along with log2 fold change, Q value, and direction of regulation (upregulated or downregulated).

**Table S1: Up and downregulated differentially expressed genes (DEGs) in female vs male Pitx2^+/+^ corneas.** This table lists the up and downregulated DEGs identified in female compared with male Pitx2⁺/⁺ corneas. For each gene, the Gene ID and gene name are provided, along with log2 fold change, Q value, and - log (Q value). Genes with a log2 fold change < −1 or > 1 are highlighted in green.

**Table S2: Gene Ontology (GO) biological process enrichment in female versus male Pitx2⁺/⁺ corneas.** This table presents GO biological process enrichment analysis comparing female and male Pitx2⁺/⁺ corneas. The table includes GO term ID, GO term description, number of candidate genes associated with each term, rich ratio, p-value, Q value, and the corresponding gene IDs and gene symbols for each enriched biological process.

**Table S3: Up and downregulated differentially expressed genes (DEGs) in female Pitx2^egl1/egl1^ vs Pitx2^+/+^ corneas.** This table lists the up and downregulated DEGs identified in female Pitx2^egl1/egl1^ compared Pitx2⁺/⁺ corneas. For each gene, the Gene ID and gene name are provided, along with log2 fold change, Q value, and - log (Q value). Genes with a log2 fold change < −1 or > 1 are highlighted in green.

**Table S4: Gene Ontology (GO) biological process enrichment in female Pitx2^egl1/egl1^ versus Pitx2⁺/⁺ corneas.** This table presents GO biological process enrichment analysis comparing female Pitx2^egl1/egl1^ and Pitx2⁺/⁺ corneas. The table includes GO term ID, GO term description, number of candidate genes associated with each term, rich ratio, p-value, Q value, and the corresponding gene IDs and gene symbols for each enriched biological process.

**Table S5: Up and downregulated differentially expressed genes (DEGs) in male Pitx2^egl1/egl1^ vs Pitx2^+/+^ corneas.** This table lists the up and downregulated DEGs identified in male Pitx2^egl1/egl1^ compared Pitx2⁺/⁺ corneas. For each gene, the Gene ID and gene name are provided, along with log2 fold change, Q value, and - log (Q value). Genes with a log2 fold change < −1 or > 1 are highlighted in green.

**Table S6: Gene Ontology (GO) biological process enrichment in male Pitx2^egl1/egl1^ versus Pitx2⁺/⁺ corneas.** This table presents GO biological process enrichment analysis comparing male Pitx2^egl1/egl1^ and Pitx2⁺/⁺ corneas. The table includes GO term ID, GO term description, number of candidate genes associated with each term, rich ratio, p-value, Q value, and the corresponding gene IDs and gene symbols for each enriched biological process.

**Table S7: Up and downregulated differentially expressed genes (DEGs) in male vs female Pitx2^+/+^ trigeminal ganglia.** This table lists the up and downregulated DEGs identified in male compared with female Pitx2⁺/⁺ trigeminal ganglia. For each gene, the Gene ID and gene name are provided, along with log2 fold change, Q value, and - log (Q value). Genes with a log2 fold change < −1 or > 1 are highlighted in green.

**Table S8: Gene Ontology (GO) biological process enrichment in male versus female Pitx2⁺/⁺ trigeminal ganglia.** This table presents GO biological process enrichment analysis comparing male and female Pitx2⁺/⁺ trigeminal ganglia. The table includes GO term ID, GO term description, number of candidate genes associated with each term, rich ratio, p-value, Q value, and the corresponding gene IDs and gene symbols for each enriched biological process.

**Table S9: Gene Ontology (GO) biological process enrichment in female Pitx2^egl1/egl1^ vs Pitx2^+/+^ trigeminal ganglia.** This table presents GO biological process enrichment analysis comparing female Pitx2^egl1/egl1^ and Pitx2⁺/⁺ trigeminal ganglia. The table includes GO term ID, GO term description, number of candidate genes associated with each term, rich ratio, p-value, Q value, and the corresponding gene IDs and gene symbols for each enriched biological process.

**Table S10: Gene Ontology (GO) biological process enrichment in male Pitx2^egl1/egl1^ vs Pitx2^+/+^ trigeminal ganglia.** This table presents GO biological process enrichment analysis comparing male Pitx2^egl1/egl1^ and Pitx2⁺/⁺trigeminal ganglia. The table includes GO term ID, GO term description, number of candidate genes associated with each term, rich ratio, p-value, Q value, and the corresponding gene IDs and gene symbols for each enriched biological process.

## Notes

### Competing Interest Statement

The authors have declared no competing interest.

